# A single neuron in *C. elegans* orchestrates multiple motor outputs through parallel modes of transmission

**DOI:** 10.1101/2023.04.02.532814

**Authors:** Yung-Chi Huang, Jinyue Luo, Wenjia Huang, Casey M. Baker, Matthew A. Gomes, Alexandra B. Byrne, Steven W. Flavell

**Affiliations:** Picower Institute for Learning & Memory, Department of Brain & Cognitive Sciences, Massachusetts Institute of Technology, Cambridge, MA, USA; Department of Neurobiology, UMass Chan Medical School, Worcester, MA, USA

## Abstract

Animals generate a wide range of highly coordinated motor outputs, which allows them to execute purposeful behaviors. Individual neuron classes in the circuits that generate behavior have a remarkable capacity for flexibility, as they exhibit multiple axonal projections, transmitter systems, and modes of neural activity. How these multi-functional properties of neurons enable the generation of highly coordinated behaviors remains unknown. Here we show that the HSN neuron in *C. elegans* evokes multiple motor programs over different timescales to enable a suite of behavioral changes during egg-laying. Using HSN activity perturbations and in vivo calcium imaging, we show that HSN acutely increases egg-laying and locomotion while also biasing the animals towards low-speed dwelling behavior over longer timescales. The acute effects of HSN on egg-laying and high-speed locomotion are mediated by separate sets of HSN transmitters and different HSN axonal projections. The long-lasting effects on dwelling are mediated by HSN release of serotonin that is taken up and re-released by NSM, another serotonergic neuron class that directly evokes dwelling. Our results show how the multi-functional properties of a single neuron allow it to induce a coordinated suite of behaviors and also reveal for the first time that neurons can borrow serotonin from one another to control behavior.

## INTRODUCTION

Individual neurons are the basic units of computation in the brain. While some neuron classes have straightforward functional roles, many others exhibit complex multi-functional roles. Single neurons can participate in multiple networks, co-release multiple neurotransmitters, and contain multiple electrotonically isolated subcellular compartments. Moreover, the diverse transmitter outputs of neurons can influence activity within downstream circuits over different timescales. In vivo studies that relate these multi-functional properties of neurons to animal behavior remain limited.

Studies across a variety of organisms have identified neuron classes with multi-functional properties. In the stomatogastric ganglion of crustaceans, individual neuron classes can be co-active with multiple oscillatory networks, each with their own rhythm, and influence activity in each of these networks (Bucher et al., 2006; Weimann and Marder, 1994). Neuron classes whose activities are associated with multiple motor programs have more recently been identified in worms, flies, and mice (Atanas et al., 2022; Fisher, 2022; Fusi et al., 2016; Ji et al., 2021). In invertebrates and vertebrates, neurons are known to co-release multiple transmitters (Svensson et al., 2018). For example, in *C. elegans*, RIM neurons control escape responses via release of glutamate, acetylcholine, tyramine, and neuropeptides (Florman and Alkema, 2022; Huo et al., 2023; Sordillo and Bargmann, 2021). In mammals, midbrain dopamine neurons co-release GABA to modulate basal ganglia circuits (Tritsch et al., 2012). Neurons with multiple electrotonically isolated compartments are also widespread in nervous systems (Hendricks et al., 2012; Stuyt et al., 2022), though the functional importance of such compartmentalization for behavioral control is not as well understood. Finally, while neurons are known to influence behavior over different timescales (Flavell et al., 2022), it is unclear whether individual neurons can influence multiple downstream behaviors over different timescales. Relating multi-functional properties of neurons to their influence on circuit activity and behavior will be critical to achieve a full understanding of the complex roles that individual neuron classes play in vivo.

The roundworm *C. elegans* provides a tractable system for dissecting the functional properties of individual neurons in the context of complex behaviors. There are exactly 302 neurons in the *C. elegans* nervous system, and the connectivity among these neurons is known (Cook et al., 2019; White et al., 1986; Witvliet et al., 2021). Each neuron in this system can be perturbed using an extensive genetic toolset. *C. elegans* exhibit a well-characterized set of motor programs (de Bono and Maricq, 2005) that are extensively coordinated with one another (Cermak et al., 2020; Collins et al., 2016; Hardaker et al., 2001; Nagy et al., 2015). These include locomotion, egg-laying, head and body posture, defecation, and feeding. Mechanisms that underlie the coordination of these motor programs are still mostly unknown.

The egg-laying behavior of *C. elegans* is controlled by a compact neural circuit located in the midbody of the animal, close to the vulval muscles whose contraction causes egg-laying (Schafer, 2006). Egg-laying is increased during exposure to food and is inhibited by aversive stimuli (Daniels et al., 2000; Fenk and de Bono, 2015; Waggoner et al., 2000; Zhang et al., 2008). A key neuron in the egg-laying circuit is the HSN neuron, which synapses onto VC neurons and vulval muscles and has a command-like role in controlling egg-laying. HSN also extends a neurite to the nerve ring in the head where it makes and receives synapses from neurons in other sensorimotor circuits. However, the function of this projection to the nerve ring remains mostly unknown. HSN releases acetylcholine, serotonin, and several neuropeptides (Schafer, 2006; Taylor et al., 2021) and receives diverse modulatory inputs (Emtage et al., 2012; Ringstad and Horvitz, 2008). In freely-moving animals, HSN exhibits calcium peaks that are sometimes accompanied by egg-laying (Collins et al., 2016; Ravi et al., 2018), and its release of serotonin and the NLP-3 neuropeptide is required for egg-laying (Brewer et al., 2019). HSN’s effects on egg-laying appear to be coordinated with changes in locomotion. Egg-laying events are commonly correlated with high-speed locomotion (Cermak et al., 2020; Hardaker et al., 2001). The AVF neuron, as well as dopamine signaling, have been implicated in the coordination of egg-laying and high-speed locomotion (Cermak et al., 2020; Hardaker et al., 2001). Somewhat paradoxically, ablation of HSN or reduced serotonin release by HSN impairs low-speed dwelling states, which are stable periods of slow locomotion on food (Aprison and Ruvinsky, 2019; Flavell et al., 2013). The cellular mechanisms that explain the diverse impacts of HSN on these distinct behaviors are still unclear.

In this study, we decipher the cellular mechanisms that allow HSN to exert multiple influences on behavior. We show that HSN plays a causal role in driving egg-laying and high-speed locomotion during egg-laying while also biasing animals towards low-speed dwelling over longer timescales. We map these behavioral effects of HSN onto different HSN transmitters and different subcellular compartments. Specifically, HSN drives an acute increase in egg-laying and high-speed locomotion via release of different sets of transmitters and distinct subcellular compartments. In addition, serotonin released from HSN acts over longer timescales to induce low-speed dwelling. This slower effect of HSN serotonin is due in part to serotonin being taken up and re-released by the serotonergic neuron NSM in the head that directly evokes dwelling. Our results reveal how a single neuron can influence a broad suite of behaviors over multiple timescales and show for the first time that neurons can ‘borrow’ serotonin from one another to control behavior.

## RESULTS

### HSN neurons evoke acute egg-laying and speeding, as well as long-term slowing

To examine the causal effect of HSN activity on different aspects of behavior (Fig. 1A), we performed a series of gain- and loss-of-function perturbations to HSN. To activate HSN, we generated a strain with HSN-specific expression of the blue light-sensitive opsin CoChR (Klapoetke et al., 2014). There were no promoters known to drive expression uniquely in HSN, so we developed an intersectional approach for cell-specific expression (Fig. 1B). We fused the *cat-4* promoter to an inverted and floxed CoChR-sl2-GFP expression cassette. Due to its inverted orientation, CoChR-sl2-GFP was not expressed in these animals. We then crossed in a transgene expressing Cre recombinase under the *egl-6* promoter, whose expression overlaps with *cat-4* exclusively in HSN. In these double transgenic animals, HSN was the only neuron that expressed GFP (Fig. S1A; we refer to this strain as HSN::CoChR). We examined the behavioral effects of activating HSN::CoChR with blue light in animals that were dwelling on a food source. As expected, this evoked an immediate increase in egg-laying (Fig. 1C). In addition, we observed a robust increase in forward locomotion speed time-locked to the light stimulus (Fig. 1C). Since HSN activation had not been shown to induce speeding before, we performed additional control experiments to ensure that this was due to activation of HSN and not another neuron with low-level CoChR expression. Specifically, we examined the effect of activating HSN::CoChR in *egl-1(gf)* mutant animals, in which the HSN neuron is specifically ablated through programmed cell death (Conradt and Horvitz, 1999). This abolished the light-induced increase in egg-laying and locomotion speed, indicating that both effects are due to HSN activation (Fig. 1D). We also examined whether the increase in locomotion speed was a consequence of egg-laying or, alternatively, a parallel output. To test this, we activated HSN in animals treated with FUDR, a DNA replication inhibitor frequently used to block germline development and sterilize animals (Mitchell et al, 1979). In these animals, there were no eggs, but HSN stimulation still acutely increased locomotion speed (Fig.1E; although HSN still elicited a robust speed increase, there was a small, but significant, deficit; also see Fig. 2 for additional mutants that uncouple HSN-induced egg-laying and speeding). This suggests that the increased locomotion is not an exclusive consequence of egg-laying mechanics or mechanosensory detection of egg-laying. Taken together, these experiments indicate that HSN activity can induce an acute increase in egg-laying and forward locomotion speed, and that the effect on locomotion speed is separable from HSN’s well-established role in controlling egg-laying.

**Figure 1.**
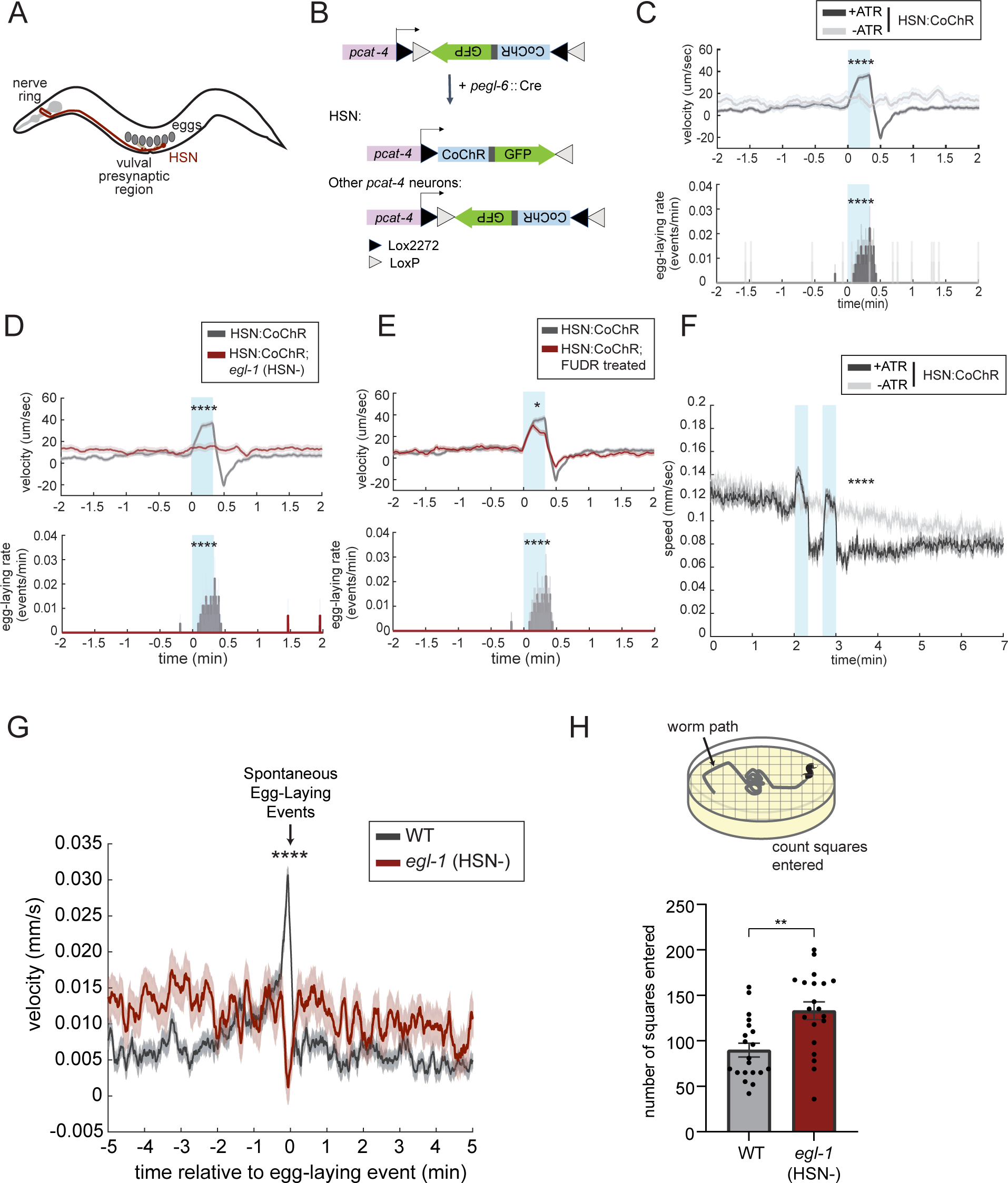
The HSN neuron evokes egg-laying, acute speeding, and long-term slowing. (A) Cartoon illustrating the anatomy of the HSN neuron (red), relative to the overall body organization of *C. elegans*. Pharyngeal and vulval muscles are shown in light gray, and eggs are shown in dark gray. (B) Cartoon depicting the intersectional promoter strategy used to obtain HSN-specific expression of CoChR. The *cat-4* promoter drives expression of an inverted CoChR-sl2-GFP expression cassette; due to its inverted orientation, it is not expressed. The *egl-6* promoter drives expression of Cre recombinase, which acts on the lox sites (black and white triangles) to invert the CoChR-sl2-GFP, permitting expression in cells that express both *cat-4* and *egl-6* (only HSN). See Fig. S1A for a fluorescent image of the resulting strain. (C) Behavioral responses to HSN::CoChR activation. Data are shown as an event-triggered average depicting the mean change in velocity (top) and egg-laying (bottom) during light illumination of HSN::CoChR animals. Blue boxes indicate the light illumination period in all HSN::CoChR experiments. There is an increase in reversals at light offset that is apparent in these data as well. We chose not to focus on this, because it was unclear if this reflected a rebound effect not directly under HSN control. Data are shown as means ± standard error of the mean (SEM). Statistics for velocity was performed on the change in velocity: difference of mean velocity during-stimulation period and velocity pre-stimulation (2-minute average baseline). Statistics for egg-laying rate was performed with the mean egg-laying rate during the stimulation period. Due to the extremely transient nature of the egg-laying events, the mean for the egg-laying rate over time is a jagged line. N = 270 stimulation events across 19 animals for the ATR group; N = 42 stimulation events across 6 animals for the no ATR control. ****p<0.0001, unpaired t-test. (D) Behavioral responses to HSN::CoChR activation in *egl-1(n487gf)* mutants, displayed as in (C). Gray shows WT behavior (same data as in C) for reference. N = 144 stimulation events across 10 animals. ****p<0.0001, unpaired t-test. (E) Behavioral responses to HSN::CoChR activation in animals treated with FUDR, which abolishes egg production. Data are displayed as in (C). Gray shows WT behavior (same data as in C) for reference. N = 140 stimulation events across 10 animals. *p<0.05, ****p<0.0001, unpaired t-test. (F) Effect of HSN::CoChR activation on speed in animals travelling at high baseline speeds, due to recent transfer to food. The mechanical agitation associated with transfer is sufficient to transiently arouse animals. The pair of two consecutive blue light stimuli were intended to mimic a burst of two HSN activity peaks (see Fig. 2), though the effects on locomotion are already visible after the first stimulus. Data are shown as speed surrounding the stimulation events. Lines depict mean speed and error shading is SEM. N= 195 animals for the ATR group, and 159 animals for the no-ATR control. Statistics was performed on the change of speed: the difference between post-stimulation (one-minute average) and pre-stimulation (one-minute average baseline). ****p<0.0001, unpaired t-test. (G) Event-triggered average showing animal speed surrounding native, spontaneous egg-laying events in wild-type (black) and *egl-1(n487gf)* (red) animals. Lines depict mean velocity and error shading is SEM. N= 389 egg-laying events across 19 animals for wild-type and 169 egg-laying events across 20 animals for *egl-1(n487gf)* animals. ****p<0.0001, unpaired t-test. (H) **Top:** Cartoon depicting the behavioral assay used to quantify exploratory behavior on food. This assay is commonly used as a metric of dwelling versus roaming behavior (Flavell et al., 2013). Animals are placed on NGM agar plates seeded with *E. coli* OP50 bacterial food as L4 animals. The next day, the tracks of the animal’s movement, which are visible in the bacterial lawn, are scored based on how many squares of a superimposed grid they traversed. **Bottom:** Exploratory behavior of animals of the indicated genotypes, using the assay depicted. Dots are individual animals, bars show means, and error bars indicate SEM. N= 20 animals for each genotype **p<0.01, unpaired t-test.

**Figure 2.**
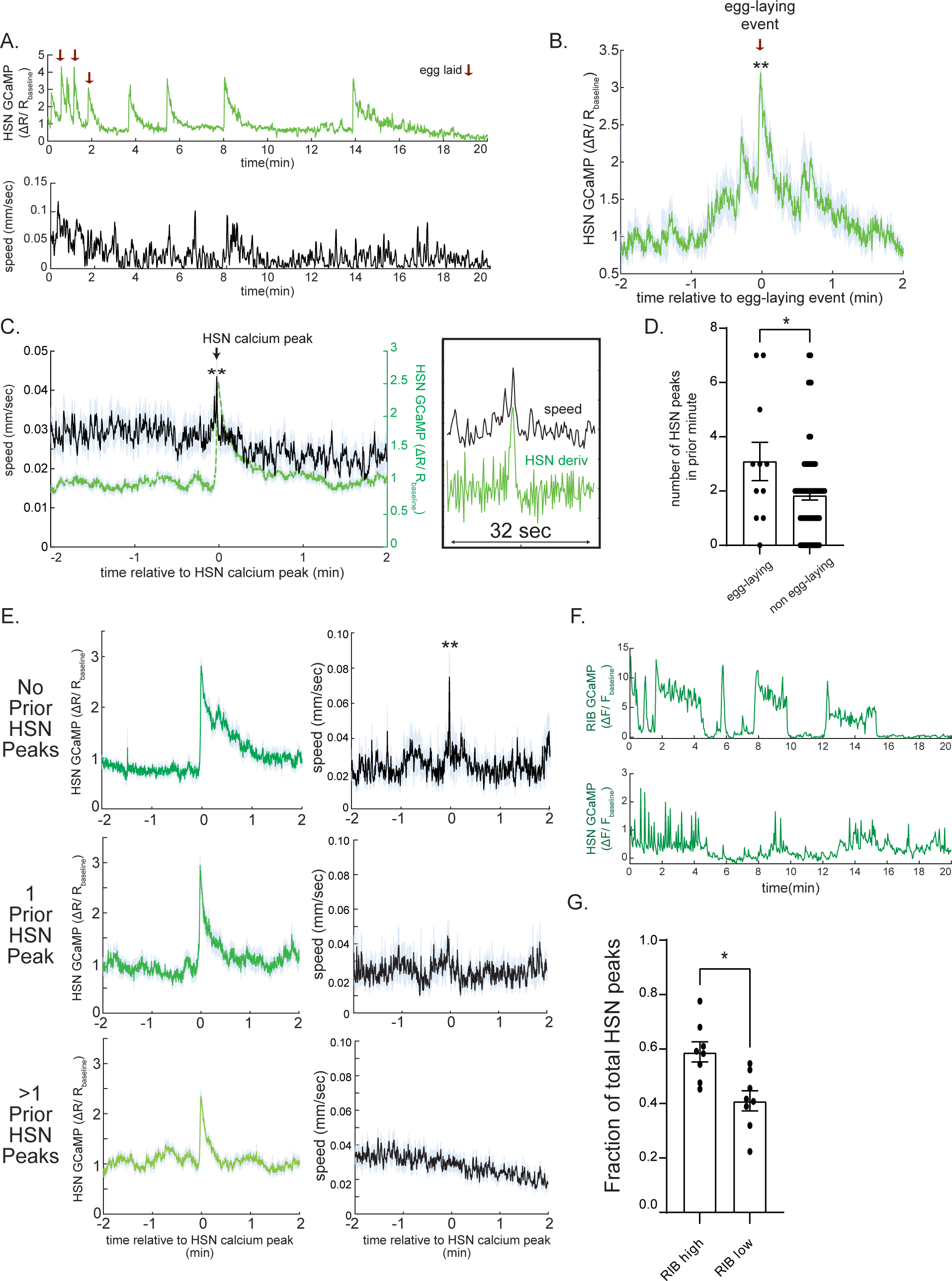
HSN activity is acutely correlated with egg-laying and high-speed locomotion. (A) Example dataset from one animal showing HSN GCaMP signal (top) and animal speed (bottom) over a 20-minute recording of HSN calcium in freely-moving animals. Red arrows indicate egg-laying events. Note that HSN activity occurs in discrete peaks and that egg-laying coincides with some of these peaks. (B) Event-triggered average showing average HSN GCaMP signal surrounding egg-laying events. Lines indicate mean and error shading is SEM. N= 16 egg-laying events. **p<0.01, empirical p-value, comparing to random distribution in Fig. S2A. (C) **Left:** Event-triggered average showing average speed (black) during HSN calcium peaks. HSN calcium is also shown (green). **Right:** the same datasets, but increased zoom along the x-axis and green here depicts the derivative of HSN GCaMP. Note the precise time alignment of animal speed and derivative of HSN calcium. Lines and error shading are means and SEM, respectively. n= 104 peaks across 15 animals. **p<0.01, increase in speed during HSN peaks, empirical p-value, comparing to random distribution in Fig. S2B. (D) Number of HSN peaks in the minute preceding HSN peaks that either result in egg-laying or not. Dots are individual HSN peaks. *p<0.05, unpaired t-test. (E) Event-triggered averages displayed as in (C), except splitting data based on how many HSN calcium peaks occurred in the minute preceding the HSN calcium peak being examined. N=22-47 calcium peaks per plot. **p<0.01, increase in speed during HSN peaks, empirical p-value, comparing to appropriate random distributions (as in (C)). (F) Example calcium traces of RIB and HSN from a joint calcium recording of both neurons in immobilized animals. (G) Fraction of the total HSN calcium peaks that occurred while RIB was high (i.e. network was in a ‘forward’ state) or low (i.e. network was in a ‘reverse’ state). Dots are individual animals. Bars show means and error bars are SEM. N= 8 animals. *p<0.05, paired t-test.

We also examined the effect of HSN activation on behavior under conditions where animals were travelling at high speed, due to recent transfer to a low-density food plate (which stimulates locomotion, unlike the conditions above where animals were already acclimated to dense food). HSN activation still caused an acute increase in locomotion during light stimulation, but this was followed by a reduction in locomotion speed that lasted for minutes (Fig. 1F; Fig. S1B). This slowing effect was most prominent when the animals were traveling at high speed and less obvious once they were acclimated on food, at which point their baseline speed was already low. This slowing effect that followed HSN activation remained intact in animals treated with FUDR, suggesting that it was not dependent on egg-laying (Fig. S1C). Together with the above results, these data suggest that HSN activation promotes acute egg-laying and high-speed locomotion, followed by a reduction in locomotion speed.

We next examined the impact of endogenous HSN activity on locomotion. For these experiments, we utilized the *egl-1(gf)* mutant animals in which HSN is specifically killed. *egl-1(gf)* mutants are known to exhibit a dramatic decrease in egg-laying (Conradt and Horvitz, 1999). We examined whether the eggs that are laid in the mutant are accompanied by an increase in locomotion speed, as is the case in wild-type animals. To do so, we recorded egg-laying and locomotion simultaneously and examined animal speed surrounding egg-laying events. Consistent with previous data (Hardaker et al., 2001), whereas wild-type animals displayed increased forward movement before and during egg-laying, this coupling was abolished in *egl-1(gf)* mutants (Fig.1G). This result indicates that HSN is required for speeding surrounding egg-laying and matches our finding that HSN activation increases speed and egg-laying. We also examined dwelling behavior in *egl-1(gf)* mutants using an assay that quantifies animal exploration across a bacterial food lawn. Indeed, as described previously, *egl-1(gf)* mutants displayed reduced dwelling behavior (or, equivalently, increased roaming; Fig.1H) (Flavell et al., 2013). This result indicates that HSN is necessary for proper dwelling and also provides a match to our findings that HSN activation can increase dwelling for minutes after optogenetic stimulation. Taken together, these data indicate that HSN is required for increased locomotion speed surrounding egg-laying events and reduced locomotion during dwelling on a bacterial food lawn.

### HSN activity is acutely correlated with egg-laying and increased locomotion

The above experiments suggest that, in addition to controlling egg-laying, HSN activity induces acute speeding and long-term slowing. Since the above experiments only examined the effects of perturbing HSN activity, we next examined how endogenous HSN activity was coupled to these behaviors in freely-moving animals. We generated a strain expressing GCaMP5A and mScarlett in HSN and performed ratiometric imaging as individual animals traveled at low speed on bacterial food lawns. Consistent with previous work, we found that HSN activity was organized into discrete calcium peaks that commonly occurred in bursts (Fig. 2A) (Collins et al., 2016; Ravi et al., 2018; Zhang et al., 2008). Across the recordings, moments of egg-laying were invariably coupled to HSN calcium peaks (Fig. 2B and S2A). However, there were also many additional calcium peaks that were not accompanied by egg-laying. To examine the relationship between HSN calcium peaks and locomotion, we examined animal speed surrounding HSN calcium peaks and found that the peaks were on average time-locked to transient increases in speed (Fig. 2C and S2B). This effect was just as robust when excluding the HSN peaks associated with egg-laying events (Fig. S2C). These data suggest that HSN calcium peaks are commonly accompanied by egg-laying and increased animal speed.

HSN calcium peaks often occurred in bursts, as previously described (Collins et al., 2016; Ravi et al., 2018; Fig. S2D shows distribution of inter-peak intervals in our data, suggesting that peaks within ~1min of one another are part of the same burst). We performed additional analyses to determine whether HSN peaks differed in their correlation with egg-laying and locomotion depending on whether the peak occurred early or later in bursts (i.e. shortly after other HSN calcium peaks). HSN calcium peaks that were associated with egg-laying most frequently occurred shortly after (<1min) several other peaks (Fig. 2D). In contrast, HSN calcium peaks were most strongly correlated with increased locomotion speed when there were no other HSN peaks in the previous minute (Fig. 2E). Thus, when several HSN calcium peaks occur in close succession, earlier peaks are more tightly associated with increased locomotion and later peaks are more associated with egg-laying. Overall, these data provide evidence that endogenous HSN activity is acutely correlated with egg-laying and speeding, matching the optogenetics results.

We next examined HSN activity in immobilized animals because it allowed us to examine the intrinsic coupling of HSN activity to motor networks without proprioceptive feedback that might couple body movement to HSN activity. In immobilized animals, the *C. elegans* nervous system exhibits robust fictive locomotion dynamics in which groups of neurons that encode forward and reverse movement switch between high and low activity states (Gordus et al., 2015; Kato et al., 2015; Uzel et al., 2022). Among these neurons, RIB is a forward-active neuron and its bi-stable activity reports the fictive locomotion state of the network. Therefore, we utilized a strain expressing GCaMP in HSN and RIB. In immobilized animals, HSN activity still occurred in discrete peaks (Fig. 2F, bottom panel). HSN calcium peaks were significantly more likely to occur when the network was in the forward, RIB-active state (Fig. 2G). These data suggest that HSN activity is coupled to the forward locomotion network state even in the absence of actual movement. Together with the freely-moving calcium imaging results above, these data suggest that HSN activity is acutely correlated with increased forward movement.

### HSN increases locomotion speed through its neuropeptidergic outputs

Having established that HSN activity is coupled to several different behavioral changes, we next sought to determine which HSN transmitter(s) mediate these effects. We first determined the transmitters that underlie acute HSN-induced speeding. To do so, we examined the effects of optogenetic HSN stimulation in mutant backgrounds lacking specific HSN transmitters. HSN evokes egg-laying via serotonin and NLP-3 neuropeptide release (Brewer et al., 2019). However, we found that animals lacking serotonin (due to a mutation in *tph-1*) or both serotonin and *nlp-3* still displayed increased speed upon HSN::CoChR activation (Fig. 3A-B; the *tph-1;nlp-3* double mutant exhibited reduced HSN-stimulated egg-laying, as expected). HSN also releases acetylcholine (Pereira et al., 2015), so we examined its impact on HSN-induced speeding. Animals deficient in acetylcholine release are uncoordinated, which makes it impossible to study HSN-induced speeding (Rand and Russell, 1984). Therefore, we used CRISPR/Cas9 genome editing to engineer a conditional knockout allele of *unc-17*, which encodes the vesicular acetylcholine transporter (VAChT) required for acetylcholine release (Fig. 3C) (Alfonso et al 1993). We first confirmed that this allele works as expected: pan-neuronal Cre expression in this strain caused animals to display an uncoordinated (Unc) phenotype. However, we found that animals with an HSN-specific deletion of *unc-17* were not uncoordinated and still displayed robust HSN-induced speeding (Fig. 3D). These data suggest that serotonin, NLP-3, and acetylcholine are not required for HSN-induced speeding.

**Figure 3.**
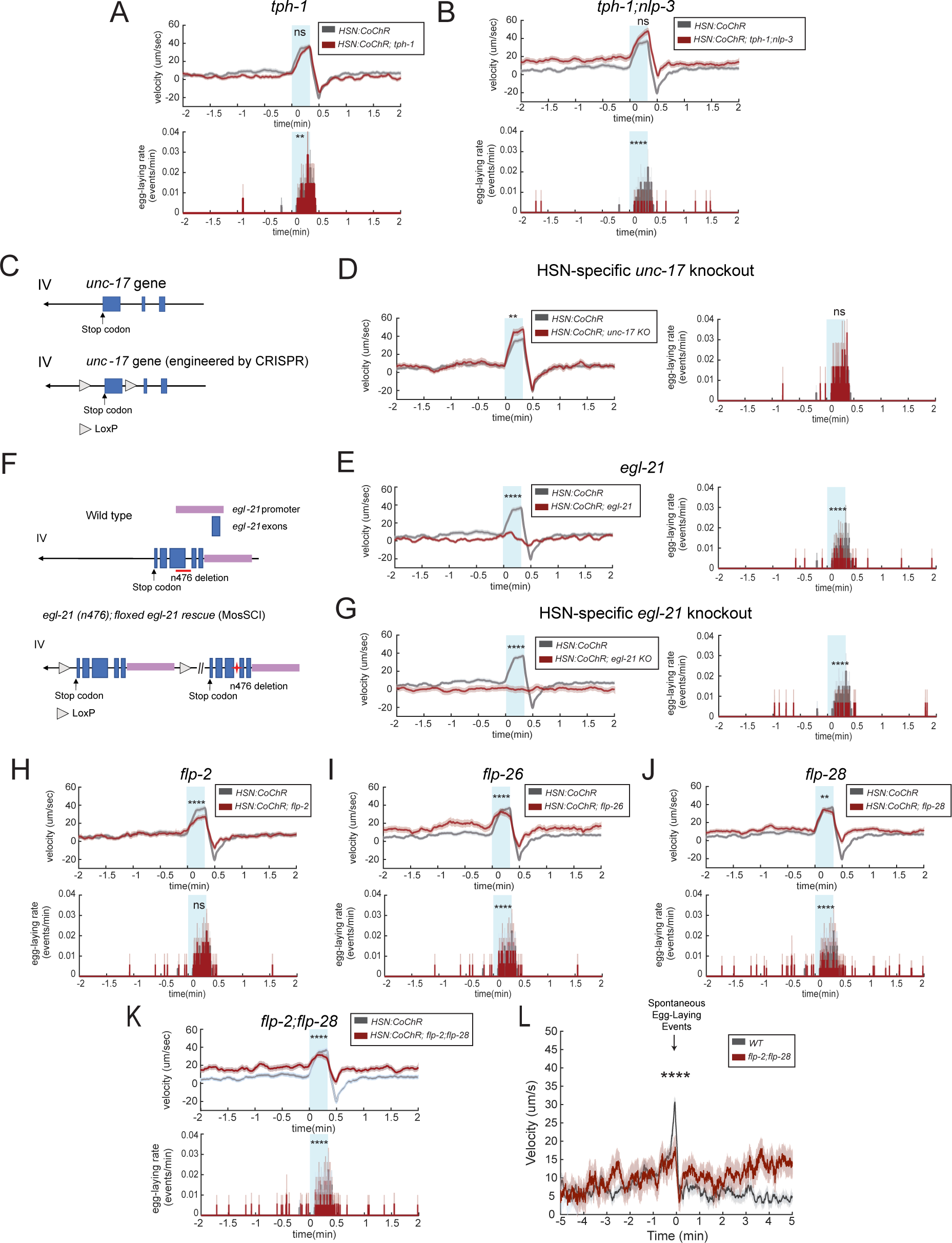
HSN evokes acute speeding through its neuropeptidergic outputs. (A-B) Event-triggered averages showing average changes in velocity (top) and egg-laying (bottom) in response to HSN::CoChR activation via light illumination. Gray provides data from WT animals as a control and reference. Mutant alleles were *tph-1(mg280)* and *nlp-3(n4897)*. Lines show means and error shading shows SEM. N= 140 stimulation events across 7 animals for *tph-1* and 180 stimulation events across 11 animals for *tph-1;nlp-3* animals. **p<0.01, ****p<0.0001, unpaired t-test. (C) Cartoon illustrating the CRISPR/Cas9-generated conditional knockout allele of *unc-17*, which encodes the vesicular acetylcholine transporter (VAChT) and is required for cholinergic transmission. (D) Event-triggered averages showing average changes in velocity and egg-laying in response to HSN::CoChR activation via light illumination. HSN-specific *unc-17* knockout refers to loxP flanked *unc-17*(*syb5779 syb5987*) animals expressing *pegl-6::Cre*. Gray provides data from WT animals as a control and reference. Lines show means and error shading shows SEM. N= 120 stimulation events across 12 animals. **p<0.01, unpaired t-test. (E) Event-triggered averages showing average changes in velocity and egg-laying in response to HSN::CoChR activation via light illumination in *egl-21* (*n476*) animals. Gray provides data from WT animals as a control and reference. Lines show means and error shading shows SEM. N=205 stimulation events across 14 animals. ****p<0.0001, unpaired t-test. (F) Cartoon illustrating the strain used for cell-specific disruption of neuropeptide production. A single-copy, floxed *egl-21* rescue was introduced into *egl-21(n476)* null animals. Red line in the top (Wild type) panel indicates the genomic location of the *n476* deletion in the genome. Red star in the lower panel indicates the *n476* mutation. (G) Event-triggered averages showing average changes in velocity and egg-laying in response to HSN::CoChR activation via light illumination. HSN-specific *egl-21* knockout refers to *egl-21(n476);kySi61[loxP-egl-21genomic-loxP]* animals expressing *pegl-6::Cre*. Gray provides data from WT animals as a control and reference. Lines show means and error shading shows SEM. N=150 stimulation events across 15 animals. ****p<0.0001, unpaired t-test. (H-K) Event-triggered averages showing average changes in velocity and egg-laying in response to HSN::CoChR activation via light illumination. Gray provides data from WT animals as a control and reference. Mutant alleles were *flp-2(gk1039)*, *flp-28(flv11)*, and *flp-26(gk3015)*, *flp-2(flv15);flp-28(flv11)* for double mutants. Because the genetic loci of *flp-2* and *flp-28* are very close, we used CRISPR to generate a 17bp insertion in *flp-2* gene in the *flp-28(flv11)* background. Lines show means and error shading shows SEM. For each genotype, *flp-2*: N= 180 stimulation events across 18 animals,; *flp-26*: 160 stimulation events across 16 animals; *flp-28*: 255 stimulation events across 26 animals; *flp-2;flp-28*: 175 stimulation events across 19 animals, **p<0.01, ****p<0.0001, unpaired t-test. (L) Event-triggered averages showing average changes in velocity surrounding native egg-laying events. Mutant alleles were *flp-2(flv15);flp-28(flv11)*,. Lines show means and error shading shows SEM. N= 167 egg laying events across 6 animals. ****p<0.0001, unpaired t-test.

Several other neuropeptide genes are also expressed in HSN, according to previous studies and genome-level expression data (Beets et al., 2022; Ripoll-Sánchez et al., 2022; Taylor et al., 2021). Thus, we next asked whether HSN neuropeptide production is required for HSN-induced speeding. To do so, we examined HSN-induced speeding in animals with a null mutation in *egl-21*, which encodes a carboxypeptidase required for processing of many neuropeptides into their mature functional forms (Jacob and Kaplan, 2003). HSN-induced speeding was abolished in these mutants (Fig. 3E). However, *egl-21* is expressed very broadly, so it was not clear whether this effect was due to loss of neuropeptide production in HSN or other neurons. Therefore, we examined HSN-induced speeding in a strain harboring an HSN-specific deletion of *egl-21*. We generated this strain by first introducing a single-copy, floxed rescue of *egl-21* into *egl-21(n476)* null mutants (Fig. 3F). We crossed HSN::CoChR into this strain. The *egl-6*::nCre transgene that acts a component of the HSN::CoChR intersectional promoter allows for HSN-specific deletion of *egl-21* (along with ~5 other neurons that express *egl-6*, such as DVA and SDQ) (Ringstad and Horvitz, 2008). Behavioral analysis of these animals revealed that they have abolished HSN-induced speeding (Fig. 3G). An analysis of baseline off-food velocity in the HSN-specific *egl-21* knockout animals confirmed that these animals were able to reach high velocity matching control animals under different environmental conditions, indicating that this deficit was not due to a general impairment in locomotion (Fig, S3A). This suggests that HSN neuropeptide production is required for HSN-induced speeding.

We attempted to determine which HSN neuropeptide(s) mediate HSN-induced speeding. To do so, we obtained a panel of 12 mutants lacking neuropeptides reported to be expressed in HSN (Beets et al., 2022; Ripoll-Sánchez et al., 2022; Taylor et al., 2021). We first examined speeding surrounding native egg-laying events for single mutants lacking these neuropeptide genes. Of these, *flp-2*, *flp-26* and *flp-28* impacted animal speed surrounding egg-laying (Fig. S3B-D; other mutants did not affect locomotion). *flp-2* has been previously shown to impact locomotion (Chen et al., 2016), whereas *flp-26* and *flp-28* have not been closely examined. To more specifically assay their involvement in HSN-induced speeding, we crossed these three mutants into the HSN::CoChR strain. We found that the loss of *flp-2*, *flp-26*, and *flp-28* all partially attenuated HSN-induced speeding, but none of these mutants had fully abolished speeding, suggesting that they may work together to drive increased forward movement downstream of HSN activation (Fig. 3H-J). Indeed, we found that HSN-induced speeding was more robustly attenuated in double mutants lacking *flp-2* and *flp-28* (Fig. 3K; binning these animals based on their pre-stimulation baseline speeds still revealed significant deficits in HSN-induced speeding, Fig. S3E-F). Correspondingly, the native coupling of egg-laying to speeding was strongly attenuated in *flp-2;flp-28* double mutants as well (Fig. 3L). Taken together, these experiments suggest that HSN acutely increases locomotion via its release of neuropeptides, including *flp-2* and *flp-28*.

### HSN induces long-term slow locomotion through the release of serotonin, which is taken up and re-released by NSM neurons

We next examined which HSN transmitter(s) drive the decrease in locomotion speed over longer time scales. For these experiments, we analyzed both baseline dwelling on food, measured through an exploration assay, and the HSN-stimulated slowing effect in which HSN::CoChR activation leads to slow locomotion for minutes afterwards (above in Fig. 1F). We found that animals lacking *tph-1* (required for serotonin synthesis) displayed decreased baseline dwelling (Fig. 4A) and a reduction in HSN-stimulated slowing (Fig. 4B). Animals lacking *nlp-3* had normal baseline dwelling (Fig. 4A) and a decrease in HSN-stimulated slowing (Fig. 4C-D). We have previously shown that cell-specific *tph-1* deletion in HSN also causes a deficit in baseline dwelling (Flavell et al., 2013). Here, we performed HSN-specific expression of *tph-1* and found that this partially rescued the *tph-1* mutant deficit in dwelling, suggesting that HSN-produced serotonin is sufficient to drive low-speed dwelling, even when other neurons do not express *tph-1* (Fig. 4E). Taken together with the above results, this suggests that HSN serotonin is not required for HSN-induced acute speeding, but is required for HSN-induced dwelling.

**Figure 4.**
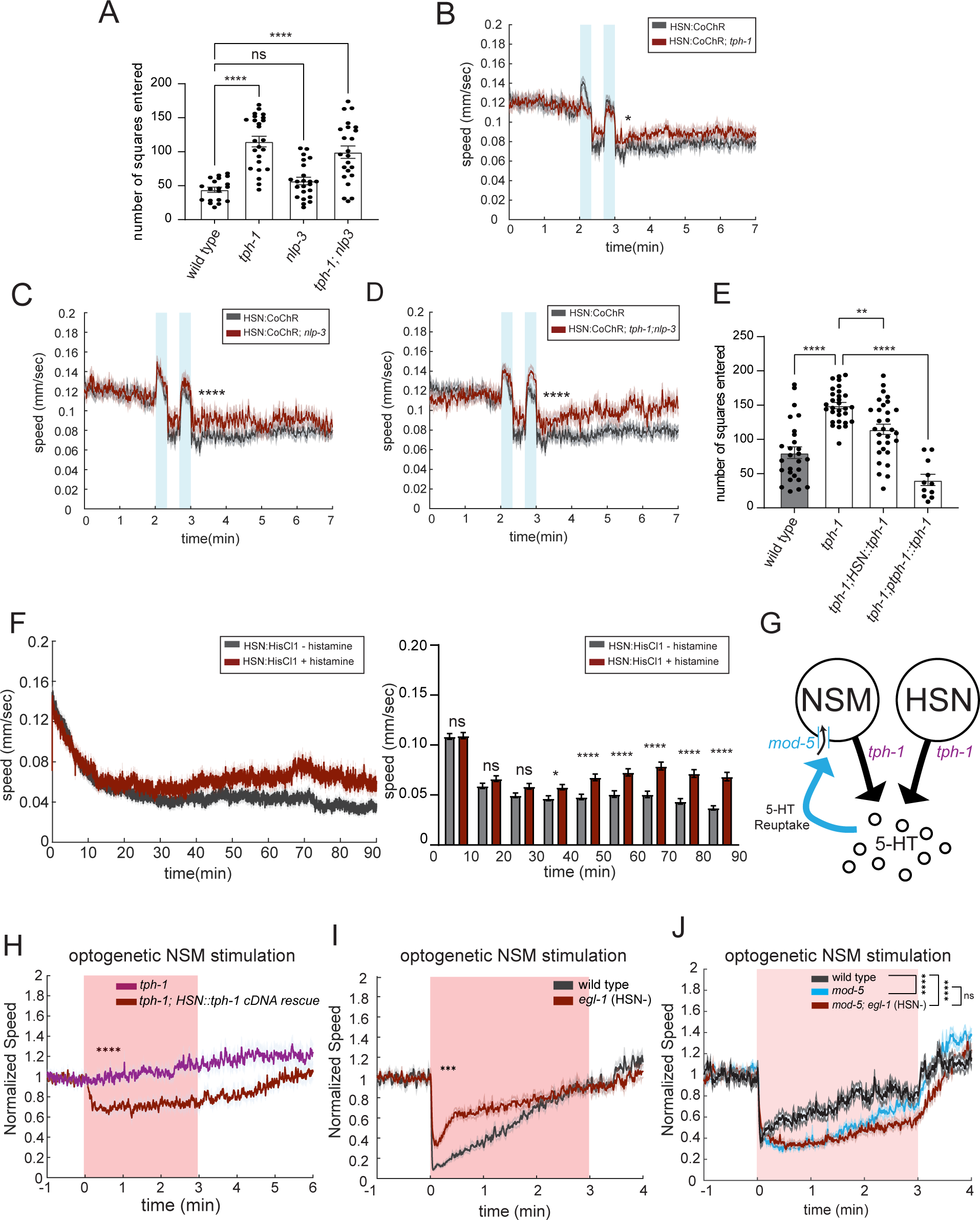
HSN-released serotonin is taken up by the NSM neuron, which re-releases it to evoke long-term slowing. (A) Exploratory behavior in animals of the indicated genotypes. Alleles used are *tph-1(mg280)* and *nlp-3(n4897)*. Dots are individual animals; bars show mean; and error bars show SEM. N= 19–23 animals for each genotype. ****p<0.0001, one-way ANOVA with Dunnett’s multiple comparison test. (B-D) Effect of HSN::CoChR activation on speed in animals travelling at high baseline speeds, due to recent transfer to food. Data are shown as mean speed surrounding light illumination for *tph-1(mg280)*, *nlp-3(n4897)*, and *tph-1(mg280);nlp-3(n4897)* animals. Lines depict mean speed and error shading is SEM. Gray provides data from WT animals as a control and reference.. N= 84 – 215 animals for each genotype. *p<0.05, ****p<0.0001, unpaired t-test. (E) Exploratory behavior in animals of the indicated genotypes. Promoters used for *tph-1* rescue are *tph-1* (3kb) promoter for native *tph-1* expressing neurons and *egl-6* promoter for HSN rescue. Dots are individual animals; bars show mean; and error bars show SEM. N= 11 – 30 animals for each genotype. **p<0.01, ****p<0.0001, one-way ANOVA with dunnett’s multiple comparison test. (F) Average speed over time of animals with HSN chemogenetically silenced (red) or not (black). HSN::HisCl refers to *pegl-6::HisCl*. Note that animals were first exposed to histamine in this experiment at t=0 min of this video. **Left:** instantaneous speed of histamine-treat and no histamine control over 90-minute recordings. **Right:** mean speed over every 10 min interval for each condition. N= 202 and 208 animals for regular NGM and histamine NGM plates respectively. *p<0.05, ****p<0.0001. Bonferroni-corrected t-test. (G) Cartoon illustrating serotonin release and re-uptake by NSM and HSN neurons. Note that HSN does not appear to express *mod-5*, based on single-cell sequence data and GFP reporter expression (Taylor et al., 2021; Duerr et al., 1999). (H) Event-triggered averages depicting the average change in animal speed upon NSM::Chrimson stimulation with red light illumination. This plot compares the effect of NSM stimulation in serotonin-null *tph-1* mutants to animals where serotonin production was restored specifically in HSN (*tph-1; egl-6::tph-1 cDNA*). Animals were starved for 3 hours before the assays, which makes the effects of serotonin on locomotion more pronounced. Lines show means; and error shading shows SEM. Statistics for velocity was performed on the average speed during light illumination period of indicated genotypes. N= 253 for *tph-1* animals and 351 for *tph-1;HSN::tph-1 cDNA* animals. ****p<0.0001, unpaired t-test. (I) Event-triggered averages depicting the average change in animal speed upon NSM::Chrimson stimulation with red light illumination in wild-type and *egl-1(n487gf)* animals. Lines show means; and error shading shows SEM. N= 105 for wild-type animals and 252 for *egl-1(n487gf)* animals. ***p<0.0001, unpaired t-test. (J) Event-triggered averages depicting the average change in animal speed upon NSM::Chrimson stimulation with red light illumination for indicated genotypes. Alleles used are *egl-1(n487gf)* and *mod-5(n822)*. Lines show means; and error shading shows SEM. N= 63-237 animals.

We next characterized the timescale over which HSN influences low-speed dwelling on food. As shown above, exogenous activation of HSN can induce low-speed dwelling with a fairly short latency (10s of seconds) and these effects last for minutes. To determine the timescale over which native HSN activity controls dwelling, we examined how long it takes for HSN silencing to alter the animal’s dwelling on food. For these experiments, we used the chemogenetic silencing tool histamine-gated chloride channel HisCl1 (Pokala et al., 2014). We first used the animal’s egg-laying behavior to characterize how quickly HSN is inactivated after HSN::HisCl animals are transferred to histamine-containing agar plates. By assaying egg-laying in a large population of animals, we found that egg-laying was inhibited within 5min of transfer, suggesting that HSN is silenced within minutes after animals are first exposed to histamine (Fig. S4A). We then determined the time course of the effect of HSN silencing on low-speed dwelling behavior. Animals were recorded right after being transferred onto histamine-containing plates (due to stimulation of animals upon transfer to new plates, both the control and histamine-treated animals have higher speed at the beginning of the video). HSN-silenced animals showed significantly higher speed only after 30-40min of exposure to histamine (Fig. 4F). This suggests that HSN needs to be inactivated for tens of minutes for there to be an increase in speed, revealing a long-lasting effect.

Given this slow time scale, we hypothesized that HSN-released serotonin may exert its influence on dwelling by contributing to the tonic pool of serotonin. In addition to directly interacting with downstream serotonin receptors, extracellular serotonin might then be taken up by other serotonergic neurons via MOD-5 (Jafari et al., 2011; Ranganathan et al., 2001), the serotonin transporter (SERT), and re-released to influence behavior (Fig. 4G and S4F). In particular, the serotonergic neuron NSM in the head expresses high levels of *mod-5* and instructively drives dwelling behavior in a serotonin-dependent manner (Rhoades et al., 2019; Sawin et al., 2000) (Fig. 4G). NSM is activated by feeding and its endogenous activity is strongly correlated with dwelling (Ji et al., 2021; Rhoades et al., 2019). Thus, we next examined whether HSN-produced serotonin could be taken up and re-released by NSM to modulate dwelling behavior.

A key prediction of this hypothesis is that NSM’s ability to induce slow locomotion via serotonin release should depend on HSN serotonin production. To test this, we used an assay where we optogenetically activated NSM. Consistent with previous work (Dag et al., 2023), we found that optogenetic NSM activation evoked robust slowing and that this effect was abolished in *tph-1* mutants (Fig. S4B). This indicates that this optogenetic assay can be used to study how NSM induces serotonin-dependent slowing. We next examined whether animals that only have *tph-1* expressed in HSN displayed a rescue in NSM-induced slowing. Indeed, we observed that restoring HSN serotonin production via HSN::*tph-1* expression partially rescued the ability of optogenetic NSM stimulation to induce slowing (Fig. 4H). This suggests that HSN serotonin production, conferred via transgenic HSN::*tph-1* expression, can partially rescue NSM-induced slowing.

We next performed a complementary experiment where we tested whether the loss of HSN in a wild-type background impairs the ability of NSM to induce slow locomotion. Here, we compared the effects of optogenetic NSM activation on locomotion in wild-type animals versus *egl-1(gf)* mutants that have HSN genetically ablated. Indeed, there was a dramatic reduction in NSM-induced slowing in *egl-1(gf)* mutants (Fig. 4I). This suggests HSN is required for normal NSM-induced slowing.

These *tph-1* rescue and HSN ablation results are consistent with two possible interpretations. First, HSN serotonin might be taken up by NSM via MOD-5/SERT, such that altering HSN serotonin production alters NSM’s ability to evoke slow locomotion via serotonin re-release. Alternatively, HSN serotonin may somehow be required downstream of NSM activation for slowing. To distinguish between these possibilities, we tested whether HSN could still influence NSM-stimulated slowing in a mutant background where *mod-5/*SERT was deleted. If the first interpretation is correct, there should be no additional effect of HSN ablation on NSM-stimulated slowing if *mod-5* is already deleted, since the *mod-5* mutation would already prevent extracellular serotonin from being taken up by NSM. Alternatively, if NSM-stimulated slowing requires a downstream function of HSN, then HSN ablation should still affect NSM-induced slowing even when *mod-5* is absent. Thus, we compared the effects of optogenetic NSM stimulation on locomotion in *mod-5* mutants versus *mod-5;egl-1(gf)* double mutants. We performed this experiment using a range of different light intensities to ensure that we were examining NSM-induced slowing under conditions where the slowing of *mod-5* animals was non-saturating (Fig. 4J; thus, note that the light stimulation intensities are lower for Fig. 4J, compared to Fig. 4I; see Fig. S4C-E for multiple light intensities). At all light levels tested, there was no difference between the two strains. This indicates that the effects of HSN on NSM-induced slowing are only observed when *mod-5*/SERT-dependent serotonin re-uptake is intact. Taken together, this set of experiments suggests that HSN serotonin is taken up and re-released by NSM to evoke slow locomotion.

### The distinct behavioral functions of HSN map onto different subcellular compartments

The above experiments suggest that HSN exerts a causal influence on behavior in at least three ways: (1) it drives acute egg-laying through serotonin and NLP-3 release; (2) it drives acute speeding through release of neuropeptides; and (3) it drives dwelling over longer time scales via serotonin release, which is taken up and re-released by NSM. We next asked how these distinct functions of HSN map onto its unique anatomy. The HSN soma is in the mid-body of the animal, just posterior to the vulva (Fig. 5A) (White et al., 1986). Each HSN neuron has a single neurite that projects into the ventral cord and then moves anteriorly towards the head. In the region where the axon passes over the vulva, HSN extends several short branches that are the sites of synapses with vulval muscles and VC neurons (we refer to this as the ‘vulval presynaptic region’) (McIntire et al., 1992, Garriga 1993). The main HSN axon continues to project anteriorly and enters the nerve ring in the head, where it forms and receives synapses with many other neurons (White et al., 1986).

**Figure 5.**
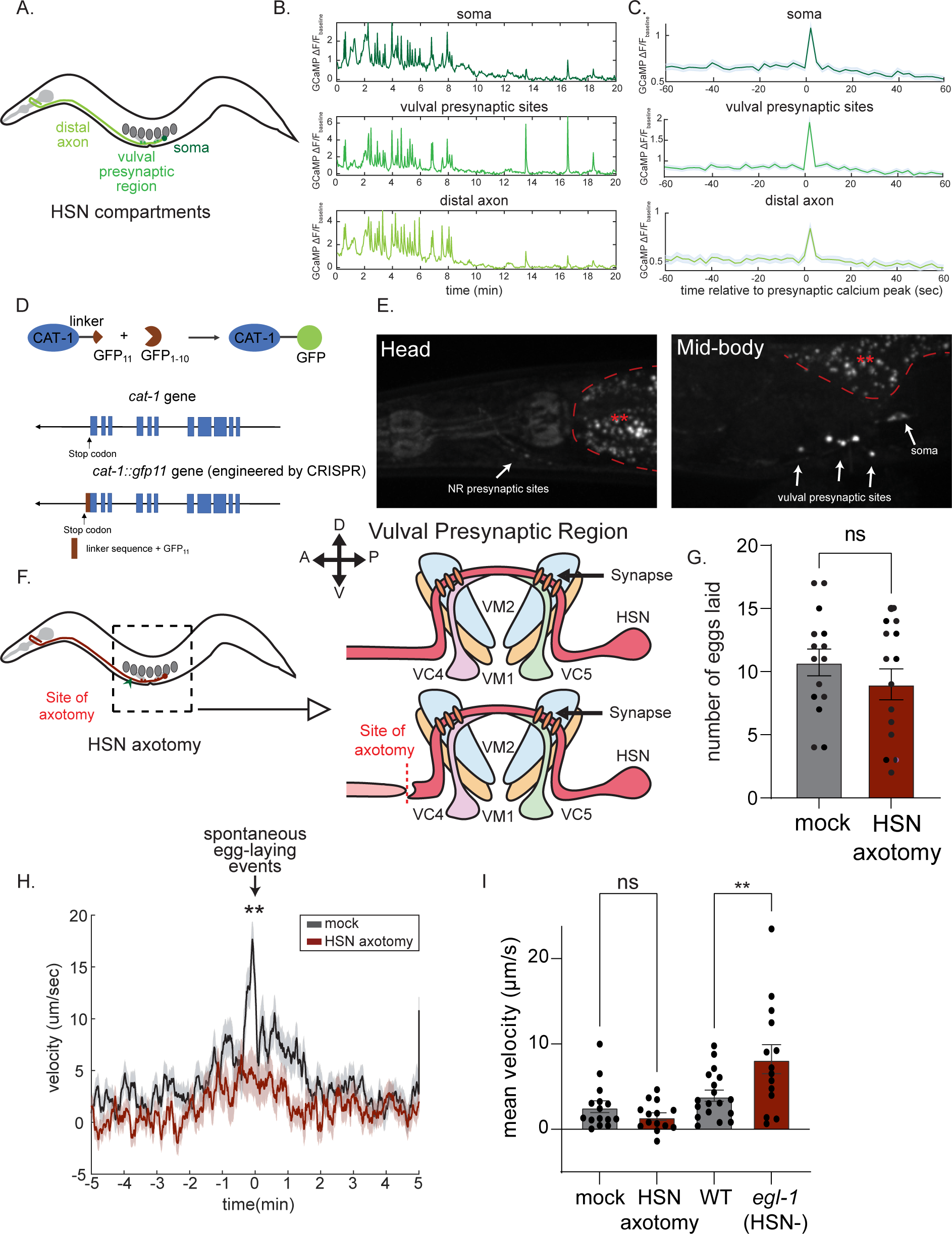
The distinct outputs of HSN map onto different sub-cellular compartments. (A) Cartoon of HSN neuron: soma, vulval presynaptic region and distal axon are labeled with different shades of green that are consistent with the colors presented in (B) and (C). (B) Example traces showing simultaneous calcium imaging of three subcellular compartments of HSN in an immobilized animal. (C) Event-triggered averages showing average HSN GCaMP signals in each sub-cellular compartment, centered on the time points of HSN calcium peaks in the vulval presynaptic region. (D) Cartoon depicting split GFP strategy to label CAT-1/VMAT in a cell-specific fashion and CRISPR/Cas9-engineered *cat-1* allele. (E) Representative images of the head (left) and mid-body (right) region of an animal with CAT-1 labeled via split GFP in HSN. Note that images were collected under identical imaging conditions and are displayed on same color scale. Soma and presynaptic sites are indicated with arrows. Areas with red asterisks are gut autofluorescence. (F) **Left:** Cartoon of HSN neuron (red) in the context of the animal’s body organization. **Right:** Illustration of the site of the HSN axotomy. Note that the HSN soma is still connected to the HSN vulval presynaptic region where HSN forms synapses onto VC neurons and vulva muscles, but not to the distal axon, as a consequence of the axotomy. (G) Egg-laying behavior of Mock and HSN-axotomized animals. Number of eggs laid over a 3hr recording is shown. ‘Mock’ animals were mounted for axotomy, but not cut with the laser, and otherwise handled identically to HSN-axotomized animals. Dots show individual animals; bars are means; and error bars are SEM. N= 15 animals for Mock and 14 animals for HSN axotomized group. (H) Event-triggered average showing average animal speed surrounding native, spontaneous egg-laying events. Data are shown for Mock and HSN-axotomized animals. Lines show means and error shading shows SEM. N= 161 egg laying events across 15 mock animals; N= 126 egg laying events across 14 HSN-axotomized animals. **p<0.01, unpaired t-test. (I) Baseline mean velocity of Mock, HSN-axotomized, wild-type and *egl-1(n487gf)* animals. Dots show individual animals; bars are means and error bars are SEM. N= 15 animals (Mock),14 animals (HSN-axotomized), 18 animals (wild-type) and 14 animals (*egl-1(n487gf))*. **p<0.01, One-way ANOVA with Sidak’s multiple comparison test.

How does HSN activity propagate across the neuron? Previous work has shown that HSN calcium peaks in the soma occur normally even when the HSN axon is severed before reaching the vulval presynaptic region (Zhang et al., 2008). This suggests that the HSN activity peaks most likely originate in the soma. To determine whether HSN soma calcium peaks are accompanied by calcium peaks along the entire HSN axon, we performed simultaneous calcium imaging of the whole HSN neuron in immobilized animals and quantified calcium transients in different regions of the neuron: the HSN soma, vulval presynaptic region, and distal axon near the nerve ring (Fig. 5B). This analysis showed that HSN calcium peaks in each compartment were accompanied by calcium peaks in the other compartments (Fig. 5C). This suggests that when HSN calcium peaks occur they are rapidly propagated across the entire neuron.

Next, we next mapped out sites of transmitter release along the HSN neurite. Here, we focused on sites of serotonin release. The *cat-1* gene encodes the *C. elegans* homolog of the vesicular monoamine transporter (VMAT) that loads serotonin into synaptic vesicles. Its subcellular localization can be used as an indicator of serotonin-containing vesicle release sites (Duerr et al, 1999). Because *cat-1* is broadly expressed, we generated a strain that allowed us to visualize endogenous CAT-1 localization in individual neurons, like HSN. We took advantage of the split GFP technique (Cabantous et al, 2005) and engineered a strain with three tandem repeats of GFP11 inserted immediately before the native *cat-1* gene’s stop codon to create an in-frame fusion protein (Fig. 5D). We then expressed GFP1-10 under the *egl-6* promoter that drives expression in HSN, but no other *cat-1*-expressing neurons. When GFP1-10 physically interacts with GFP11 it reconstitutes a functional GFP fluorophore. This approach allowed us to visualize endogenous CAT-1-containing serotonin release sites specifically in HSN. CAT-1::GFP in HSN displayed punctate localization, suggestive of presynaptic release sites. Fluorescent puncta were brightest in the vulval presynaptic region, with much weaker punctate fluorescence in the head and no expression along the neurite (Fig. 5E). This suggests that HSN primarily releases serotonin in the vulval presynaptic region.

We also tested whether the different behavioral functions of HSN require its axonal projection to the nerve ring. To do so, we examined the behavior of animals in which we specifically axotomized the HSNL/R neurites just anterior to the vulval presynaptic region in a location where the neurites were still defasciculated from the ventral nerve cord (Fig. 5F). This chosen axotomy site leaves the HSN soma connected to the vulval presynaptic region but not to the remainder of the neurite that projects into the nerve ring. We used a pulsed nitrogen laser to cut both HSN neurites in this position in L4 animals (Fig. 5F). Animals were then recovered and after 24 hours their behavior was compared to ‘mock’ animals that were immobilized and recovered in the same manner, but were not axotomized. We found that egg-laying rates were unaffected by laser axotomy at this position, consistent with the notion that the HSN vulval presynaptic region, not the projection to the head, controls egg-laying (Fig. 5G). However, we found that egg-laying events in the axotomized animals displayed much weaker coupling to increased locomotion speed (Fig. 5H). This suggests that HSN signal propagation to the head is required for proper speeding during egg-laying. Finally, we found that the baseline speed of animals on food was not significantly affected by HSN axotomy (Fig. 5I). Thus, disrupting the HSN axonal projection to the nerve ring does not bias the animals towards roaming, even though HSN cell ablation or HSN silencing does (Fig. 5I; the effect of *egl-1(gf)*) is also shown here for reference). This suggests that HSN signal propagation to the head is not necessary for the dwelling-promoting function of HSN. This is consistent with our observation that most HSN serotonin release, which promotes dwelling, occurs in the vulval presynaptic region. Altogether these experiments suggest that HSN axonal projections to the head are required for acute speeding during egg-laying but not baseline egg-laying or on-food locomotion.

### Sensory control of egg-laying is mediated by humoral release of neuropeptides by sensory neurons

The above results above provide information about how the functional outputs of HSN map onto the different anatomical and molecular features of this neuron. We also wanted to examine how sensory inputs are transmitted to HSN, given its unique anatomy and function. Egg-laying behavior is impacted by many aspects of the sensory environment, including food availability, aversive cues, and more (Fenk and de Bono, 2015; Liu et al., 2018; Teshiba et al., 2016; Trent et al., 1983). Here, we focused on the effects of osmolarity as it has been shown that a mild upshift in osmolarity (< 1 Osm) triggers aversive behavioral responses, including reduced egg-laying (Yu et al., 2017; Zhang et al., 2008). Previous work has also shown that HSN calcium peaks are inhibited by high osmolarity (Zhang et al., 2008), suggesting that this may be a good system for studying sensory control over HSN activity and behavior.

To examine how high osmolarity inhibits HSN and egg-laying, we used a relatively high-throughput behavioral assay where animals were transferred to a high osmolarity plate for one hour and the number of eggs laid was counted (Fig. 6A). Standard *C. elegans* growth plates in the lab are 150 mOsm (Brenner, 1974). To increase osmolarity, we added sorbitol at defined concentrations up to 450 mOsm. This revealed a dose-dependent effect whereby higher levels of osmolarity in the plate inhibited egg-laying (Fig. 6B). In principle, this effect could be mediated by sensory detection of external or internal osmolarity levels. To determine the sensory mechanisms that link osmolarity to egg-laying, we first examined the behavioral responses of animals lacking either *tax-2* or *ocr-2*, which encode ion channels required for sensory transduction in different sets of *C. elegans* sensory neurons (Ferkey et al., 2021). While *ocr-2* mutants still responded to the osmotic stimulus, we found that the *tax-2* mutation attenuated the effect of high osmolarity on egg-laying for the 300 mOsm exposure (Fig. 6B). Higher concentrations still inhibited egg-laying in *tax-2*, suggesting that additional mechanisms may inhibit egg-laying under very high osmolarity conditions. We also observed the same behavioral phenotype in animals lacking *tax-4*, which encodes ion channel subunits that function together with those encoded by *tax-2* (Fig. 6C) (Ferkey et al., 2021). This suggests that mild increases in osmolarity inhibit egg-laying through a mechanism that involves *tax-2/4*-mediated sensory transduction.

**Figure 6.**
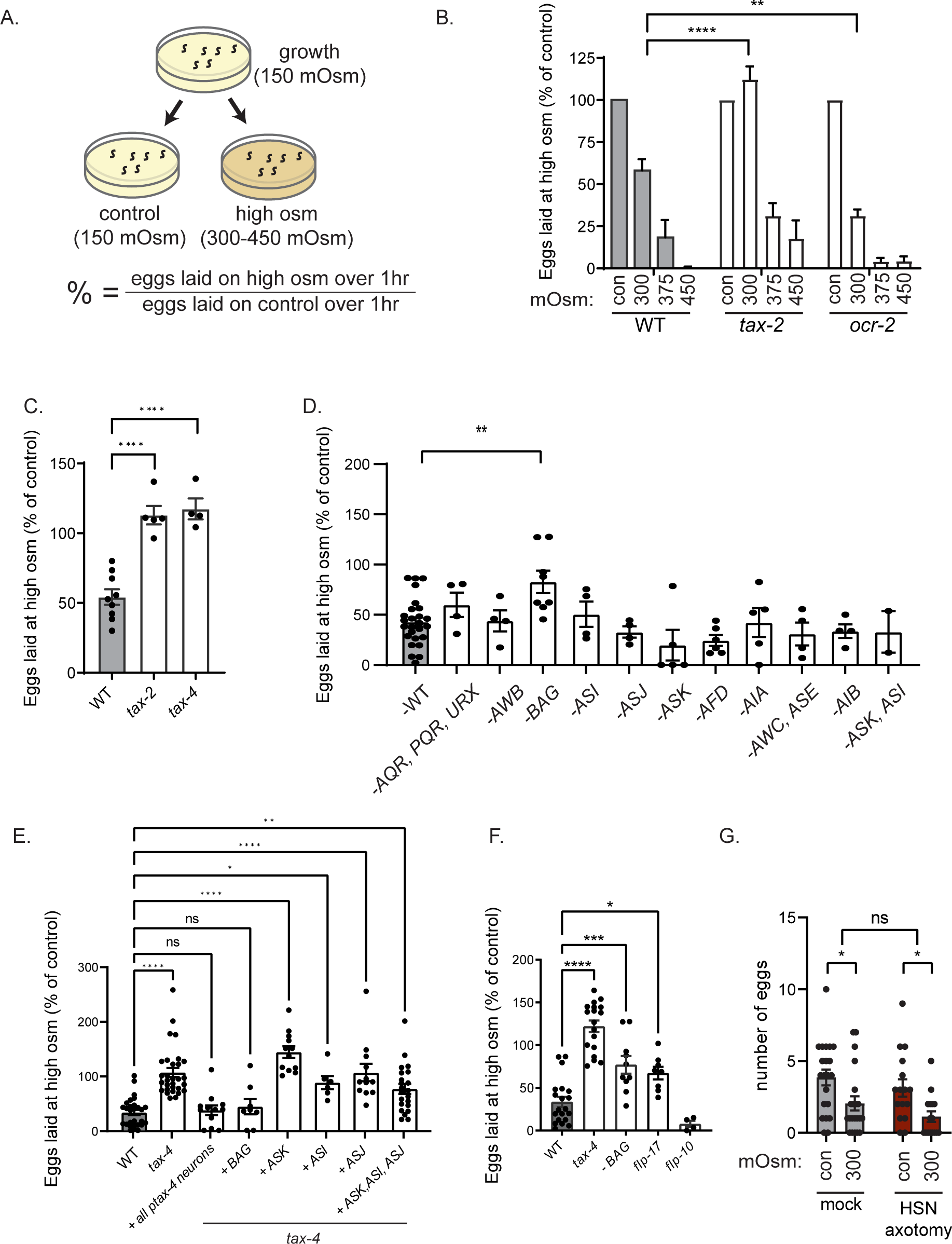
Aversive sensory input to HSN is transmitted through humoral neuropeptide release, which inactivates HSN. (A) Cartoon depicting the behavioral assay used to measure the effect of high osmolarity on egg-laying behavior of *C. elegans*. The metric at the bottom is the y-axis in subsequent plots. (B) Percent eggs laid on high osmolarity, compared to control (150 mOsm) osmolarity, shown for the indicated genotypes. Alleles used are *tax-2(p691)* and *ocr-2 (ak47)*. N= 3 – 4 plates with 10 animals on each plate. Bars are means and error bars are SEM. **p<0.01, ****p<0.0001, two-way ANOVA with dunnett’s multiple comparison test. (C) Percent eggs laid on high osmolarity (300 mOsm), compared to control (150 mOsm) osmolarity, shown for the indicated genotypes. Alleles used are *tax-2(p691)and tax-4 (p678)*. Dots are individual plates with 10 animals each; bars are means; and error bars are SEM. N= 4 – 9 plates for each genotype.. ****p<0.0001, one-way ANOVA with dunnett’s multiple comparison test. (D) Percent eggs laid on high osmolarity (300 mOsm), compared to control (150 mOsm) osmolarity, shown for the indicated genotypes. Dots are individual plates with 10 animals each; bars are means; and error bars are SEM. N= 2 – 26 plates for each genotype. Exact ablation lines used are: **AWB**: *peIs1715 [str-1p::mCasp-1 + unc-122p::GFP]*; **AQR/PQR/URX:** *qals2241[gcy-36::egl-1; gcy-35::GFP; lin-15(+)]*; **BAG:** *kyIs536 (flp-17p::p17 domain of human Caspase3::sl2::gfp; elt-2::GFP); kyIs538 (glb-5p::p12 domain of human Caspase3::sl2::gfp; elt-2::mcherry)*; **ASI:** *oyIs84(gpa-4p::TU#813+gcy-27p::TU#814+gcy-27p::GFP+unc-122p::DsRed) TU#813 and TU#814 are split caspase vector*; **ASJ:** *mgIs40([daf-28p::nls-GFP]; jxEx100[trx-1::ICE + ofm-1::gfp])*; **ASK:** *qrIs2[sra-9::mCasp1]*; **AFD**: *ttx-1(p767)*; **AIA**: *kyEX4745[gcy-28dp::unc-103(gf)::sl2::mCherry, elt-2::mCherry]*; **AWC, ASE:** *ceh-36(ky640)*; **AIB:** *flvEx356[inx-1::unc-103::sl2GFP; myo-2::mCherry]*: **ASK, ASI:** *oyIs84(gpa-4p::TU#813+gcy-27p::TU#814+gcy-27p::GFP+unc-122p::DsRed); qrIs2[sra-9::mCasp1]*. **p<0.01, one-way ANOVA with dunnett’s multiple comparison test. (E) Percent eggs laid on high osmolarity (300 mOsm), compared to control (150 mOsm) osmolarity, shown for the indicated genotypes. Dots are individual plates with 10 animals each; bars are means; and error bars are SEM. N= 6 – 28 plates for each genotype. Promoters used for *tax-4* rescue are: *tax-4* (all *tax-4* expressing neurons), *gcy-33* (BAG), *sra-9* (ASK), *srg-47* (ASI), *srh-11* (ASJ). *p<0.05, **p<0.01, ****p<0.0001, one-way ANOVA with dunnett’s multiple comparison test. (F) Percent eggs laid on high osmolarity (300 mOsm), compared to control (150 mOsm) osmolarity, shown for the indicated genotypes. Dots are individual plates with 10 animals each; bars are means; and error bars are SEM. N= 4 – 20 plates for each genotype. Alleles used are *tax-4(p678)*, BAG ablation (*kyIs536 (flp-17p::p17 domain of human Caspase3::sl2::gfp; elt-2::GFP); kyIs538 (glb-5p::p12 domain of human Caspase3::sl2::gfp; elt-2::mcherry)*), *flp-17* (*n4894*), *flp-10* (*ok2624*). *p<0.05, **p<0.01, ****p<0.0001, one-way ANOVA with dunnett’s multiple comparison test. (G) Number of eggs laid on control (150 mOsm) and high (300 mOsm) osmolarity, shown for Mock and HSN-axotomized animals (same site of HSN axotomy as shown in Fig. 5F). Dots show individual animals; bars are means; and error bars are SEM. N= 21 animals for Mock and 16 animals for HSN axotomized group. *p<0.05, two-way ANOVA followed by Sidak’s multiple comparison test.

To map out which exact sensory neurons are required for osmolarity-induced egg-laying inhibition, we performed two sets of experiments. First, we examined behavioral responses in a panel of transgenic and mutant strains with specific sensory neurons ablated. Second, we examined behavioral responses in animals that had *tax-4* rescued in different subsets of sensory neurons. We examined the effects of cell ablation for >10 sensory cell types and only observed an attenuated behavioral response in animals with the sensory neuron BAG ablated (Fig. 6D). We examined the effects of restoring *tax-4* expression in seven neuron classes and observed the most robust rescue when *tax-4* was rescued in BAG (Fig. 6E; matching the full rescue when *tax-4* was expressed under its own promoter). These results suggest that sensory transduction in BAG sensory neurons is important for osmolarity-induced inhibition of egg-laying. Interestingly, BAG is known to sense gases and its ciliated sensory ending is inside the animal’s body cavity, rather than the external environment (Ferkey et al., 2021; Witham et al., 2016). This raises the possibility that BAG may sense the animal’s internal osmolarity and, in turn, relay this information to the egg-laying circuit.

We attempted to characterize the BAG signal that impacts HSN activity and egg-laying. BAG is known to release *flp-17* and *flp-10* neuropeptides, which both function to inhibit HSN, so we examined osmolarity-induced behavioral responses in mutants lacking these neuropeptides (Ringstad and Horvitz, 2008). Indeed, *flp-17* mutants showed an attenuated egg-laying behavioral response, matching the BAG-ablated animals (Fig. 6F). This suggests that *flp-17* is important for osmolarity-induced egg-laying inhibition. Related to this, we also found that HSN axotomy (in the same position described above) did not attenuate the effects of osmolarity on egg-laying (Fig. 6G). This suggests that a humoral signal is relayed from sensory neurons to HSN to inhibit egg-laying, rather than local synaptic signaling in the head. Overall, these results are suggestive that increased osmolarity activates a BAG-FLP-17 humoral signal to inhibit HSN and egg-laying.

## DISCUSSION

Animals exhibit a wide range of motor outputs that are extensively coordinated with one another as they unfold over time. Here, we show how specific multi-functional properties of the HSN neuron endow this cell with the ability to orchestrate a suite of behavioral changes. Through causal perturbations to HSN, as well as in vivo calcium imaging, we found that HSN promotes an acute increase in egg-laying and high-speed movement, followed by low-speed dwelling behavior. We then mapped these effects onto different HSN transmitter systems and subcellular compartments. HSN’s acute effects on egg-laying and speeding are mediated by distinct sets of HSN transmitters and different subcellular compartments where HSN forms synapses with different cells. The longer lasting effect of HSN on dwelling behavior is mediated by its release of serotonin, which is taken up and re-released by serotonergic NSM neurons that directly evoke dwelling. Our results illustrate how cellular morphology, multiple transmitter systems, and non-canonical modes of transmission like neurotransmitter “borrowing” ultimately endow a single neuron with the ability to orchestrate multiple features of a behavioral program.

While the role of HSN in egg-laying is well-established (Collins et al., 2016; Conradt and Horvitz, 1999), we used a new set of tools to perturb and measure HSN activity, which revealed novel behavioral functions. First, HSN releases multiple neuropeptides to acutely increase locomotion speed prior to egg-laying. While a role for HSN neuropeptides in triggering a fast behavioral change at first seems non-traditional, our axotomy data suggest that the speed-evoking effect of HSN requires its axon in the head. Given that the locomotion circuit is located in the head and that HSN calcium peaks are reliably transmitted along the HSN axon to the head, these peptides may be well positioned to exert fast, direct action on locomotion circuits during HSN calcium peaks. We also found that HSN can induce dwelling behavior that lasts for minutes after HSN activity ends and that HSN-released serotonin was critically required for this function. Interestingly, we found that HSN-released serotonin is taken up and re-released by the NSM neuron in the head, which directly evokes dwelling. Previous work has shown that once serotonin has been released, it can be absorbed by diverse cell types (Ghai et al., 2009; Jafari et al., 2011; Lebrand et al., 1996). In *C. elegans*, these cell types include NSM, but also AIM and RIH neurons (Jafari et al., 2011; Ranganathan et al., 2001; Zhang et al., 2005). However, it has remained unclear whether this absorption is simply for serotonin turnover/degradation or, alternatively, whether this serotonin is re-released to impact circuit activity and behavior. Our work here provides direct evidence that serotonin can be transferred between serotonergic neurons and re-released to control behavior. This mechanism of neurotransmitter “borrowing” may be functionally important. NSM is directly activated by food ingestion and its release of serotonin evokes dwelling (Dag et al., 2023; Rhoades et al., 2019). Our results here suggest that NSM’s serotonin levels are influenced by HSN activity. This would then allow HSN activity to have a priming-like effect, where its recent activity could influence NSM-stimulated slowing driven by food ingestion. Future studies could make use of fluorescent serotonin sensors to define the precise spatiotemporal dynamics of extra-synaptic serotonin in *C. elegans* and investigate the mechanisms that control these dynamics.

The behavioral coordination that HSN facilitates during egg-laying has the potential to be evolutionarily adaptive. One possible reason that animals may display increased locomotion right at the moment of egg-laying is that this may deform and depolarize muscle cells adjacent to vulval muscles, facilitating the egg-laying process. Separate from this, high-speed movement during egg-laying may also allow animals to distribute their eggs in a favorable environment rather than depositing them all in one location. Such bet-hedging strategies can have adaptive advantages. Coupled with a long-term bias towards dwelling induced by the slower effects of HSN serotonin, this suite of behavioral changes could allow animals to distribute their eggs within a high-quality environment for their offspring and spend more time dwelling in the overall vicinity as well. *C. elegans* egg-laying is impacted by many aspects of the sensory environment – food, aversive cues, and more (Schafer, 2006). We found that high osmolarity, an aversive stimulus, signals through a humoral factor to alter egg-laying. HSN may integrate this aversive information with other sensory information to induce egg-laying and its associated behavioral program in maximally favorable environments. Future work in more complex sensory environments could further elucidate such integration mechanisms. Producing offspring in a favorable environment is a crucial evolutionary decision, so having a single, command-like neuron that controls this decision and coordinates it with other behaviors may be critical for fitness. Indeed, command-like neurons for egg-laying have been found in other animals like *Drosophila* as well (Vijayan et al., 2021; Wang et al., 2020). Our findings here reveal how specific multi-functional properties of such neurons can in fact allow them to induce entire suites of behavioral changes that unfold over multiple timescales.

## ACKNOWLEDGMENTS

We thank Mark Alkema, Cori Bargmann, and members of the Flavell lab for critical reading of the manuscript. We thank Cori Bargmann for guidance during mosSCI strain generation. We thank the Caenorhabditis Genetics Center (supported by P40 OD010440), Bob Horvitz, Piali Sengupta, and Cori Bargmann for strains. We thank Yun Zhang and Nuria Flames for sharing plasmids and DNA sequences. A.B.B. acknowledges funding from NIH (NS124338). Y-C.H. was supported by a Picower Fellowship. S.W.F. acknowledges funding from NIH (GM135413); NSF (Award #1845663); the McKnight Foundation; Alfred P. Sloan Foundation; The Picower Institute for Learning and Memory; and The JPB Foundation.

## AUTHOR CONTRIBUTIONS

Conceptualization, Y-C.H. and S.W.F. Methodology, Y-C.H., J.L., W.H., C.M.B., M.A.G., A.B.B., and S.W.F. Software, Y-C.H., C.M.B., and S.W.F. Formal analysis, Y-C.H. and S.W.F. Investigation, Y-C.H., J.L., W.H., C.M.B., M.A.G., A.B.B., and S.W.F. Writing – Original Draft, Y-C.H. and S.W.F Writing – Review & Editing, Y-C.H. and S.W.F. Funding Acquisition, A.B.B. and S.W.F.

## DECLARATION OF INTERESTS

The authors have no competing interests to declare.

## MATERIALS AND METHODS

### Key Resources Table

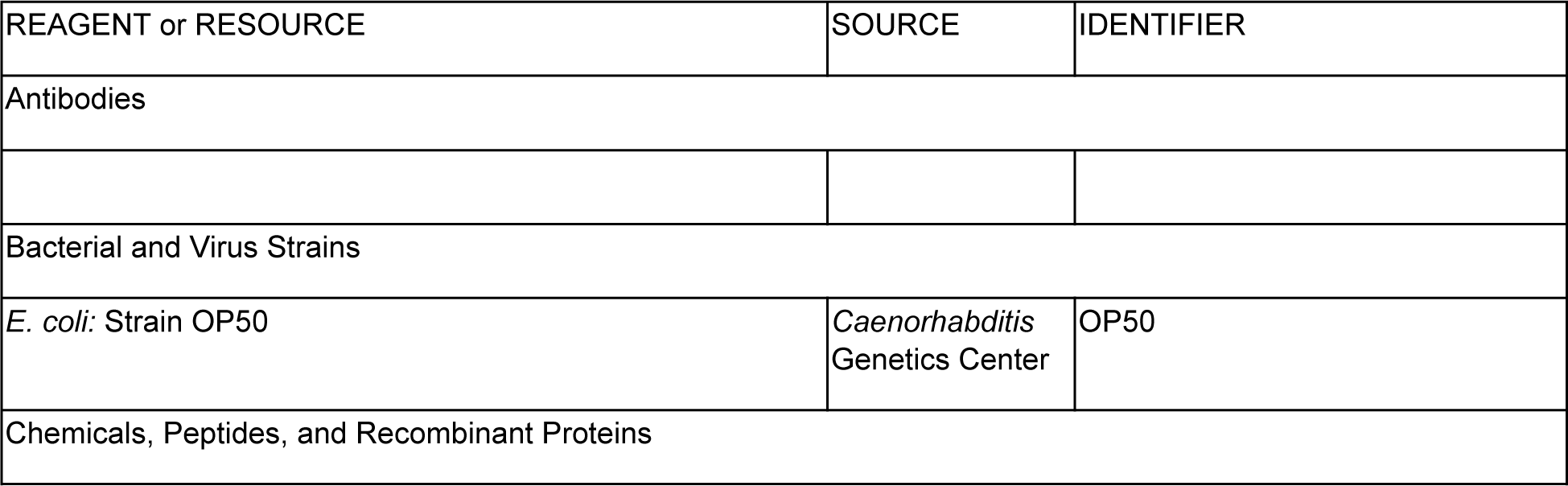
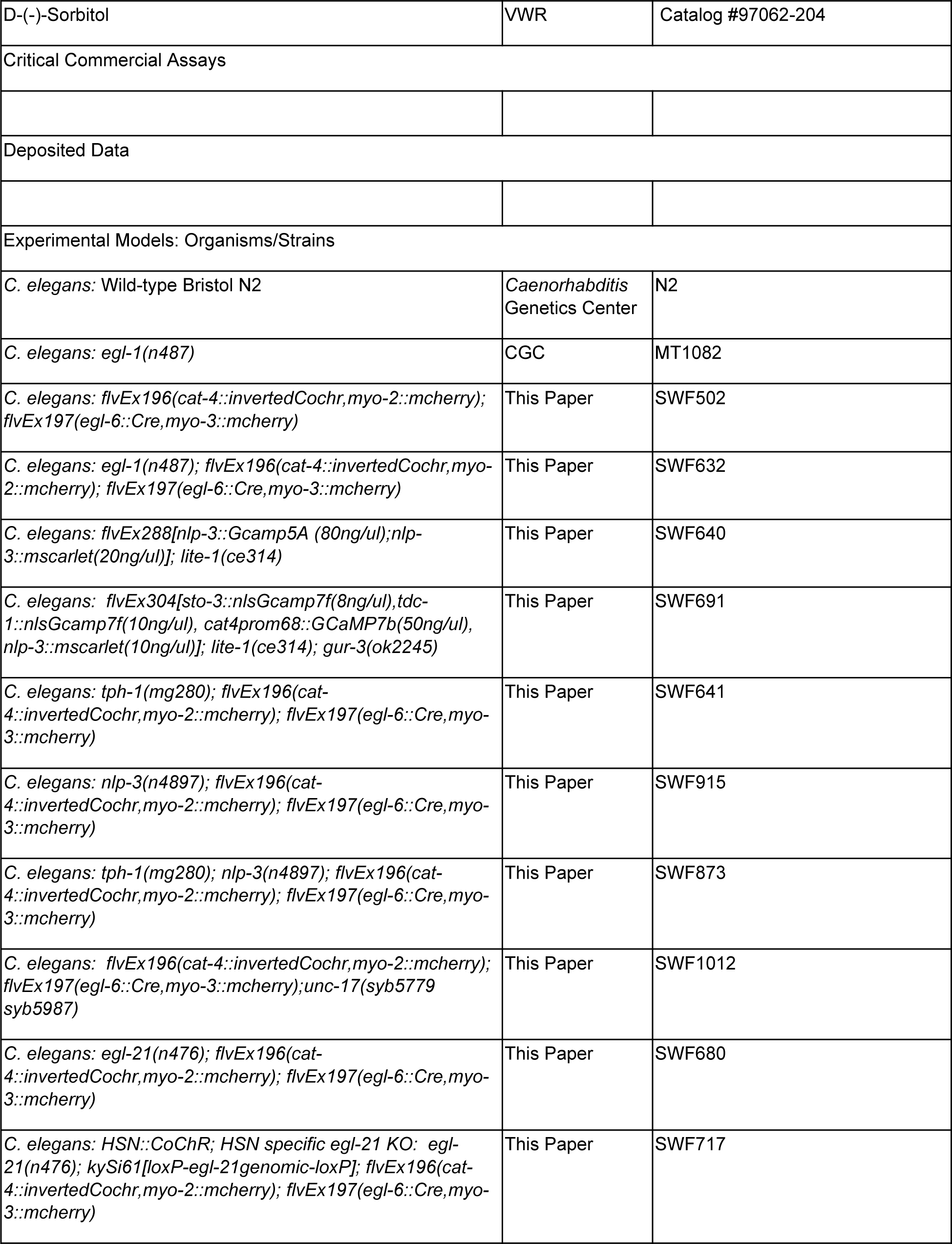
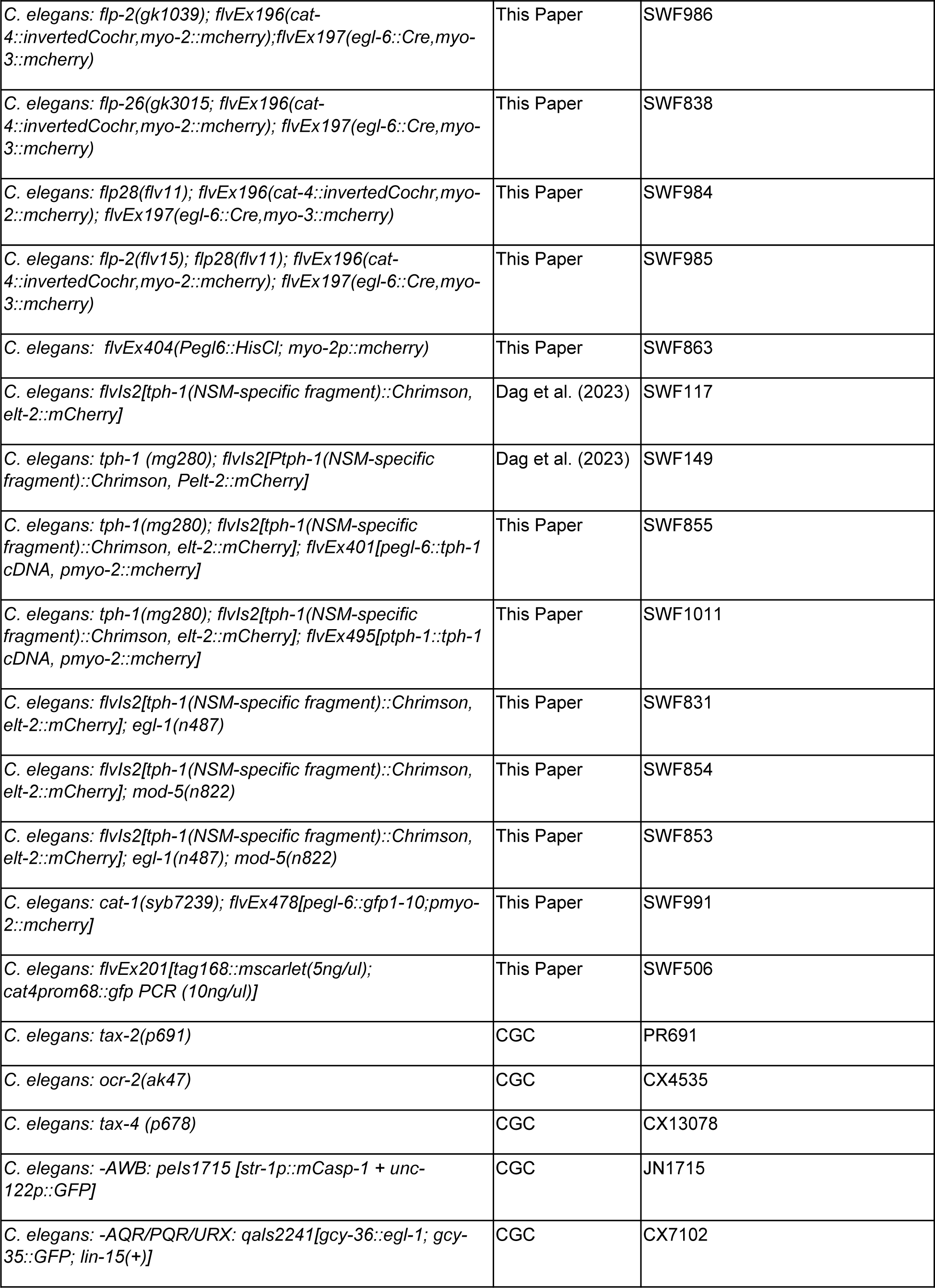
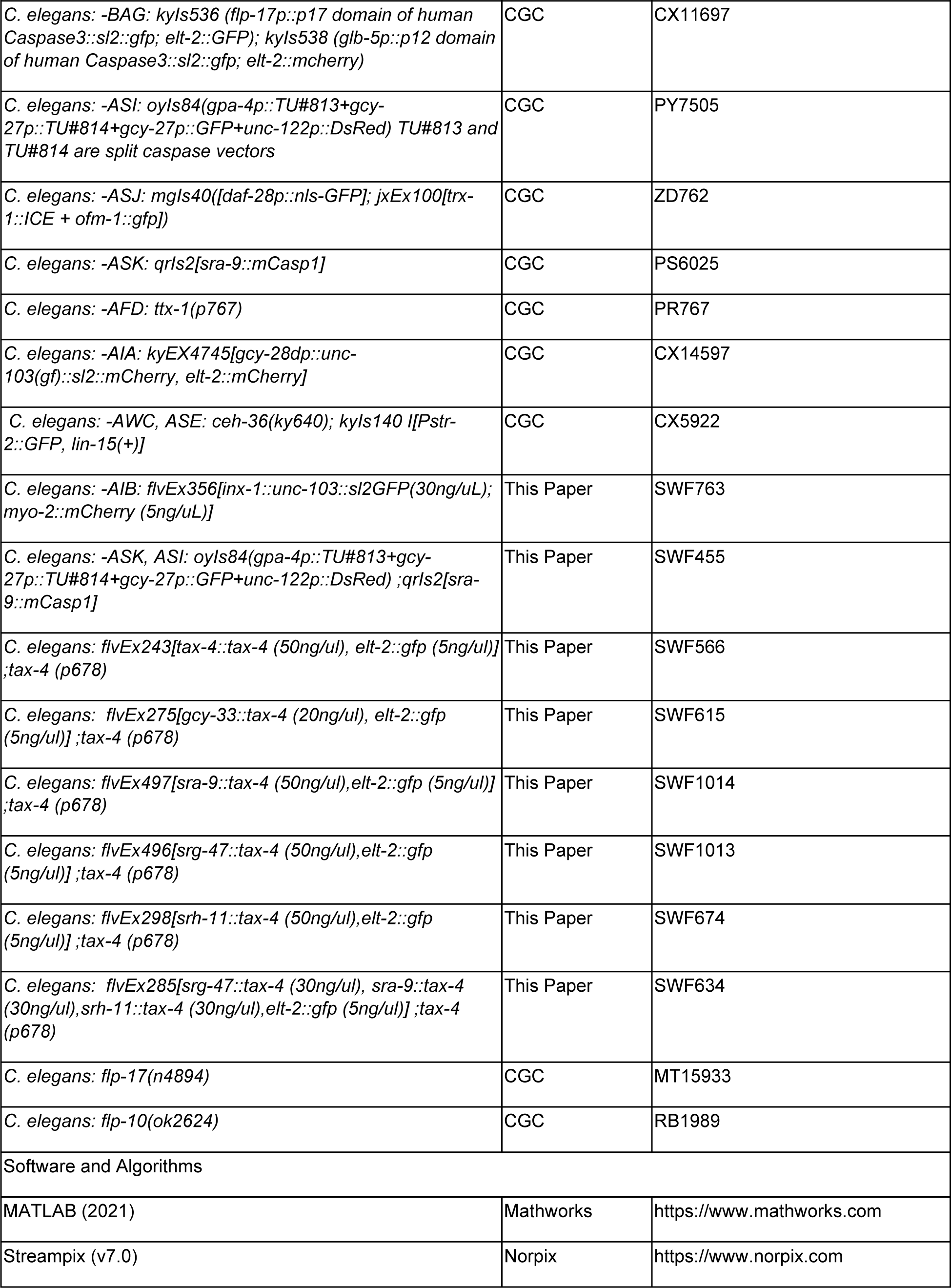
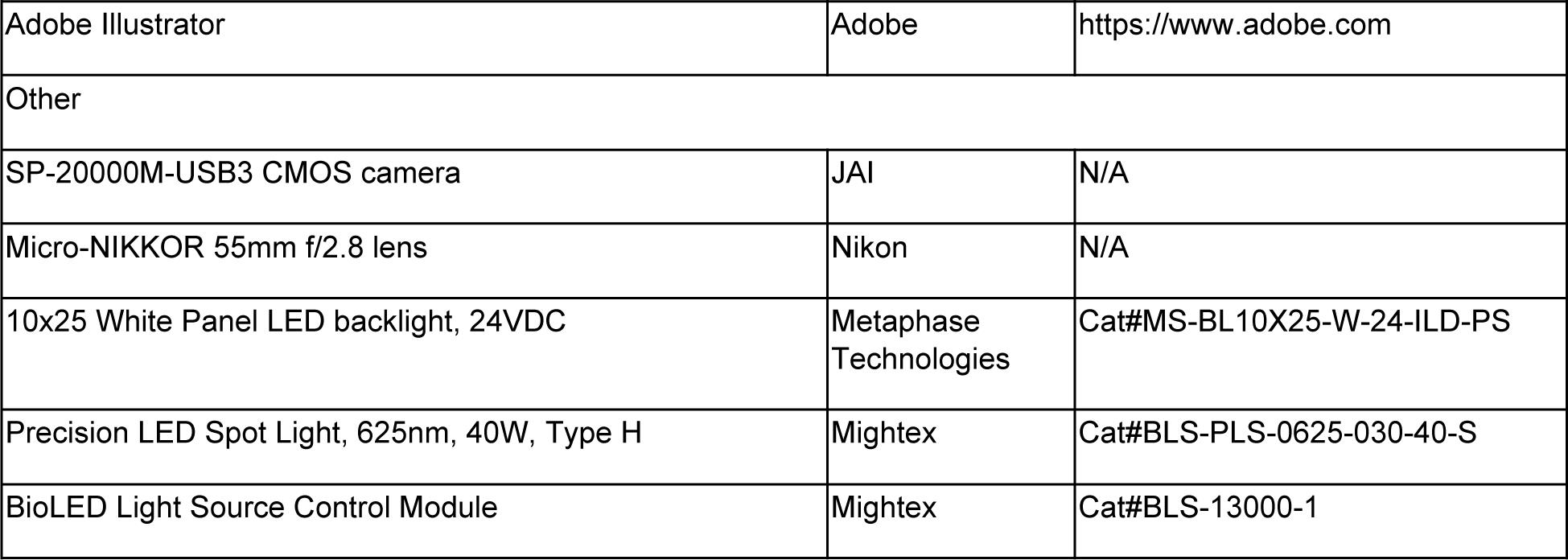

| REAGENT or RESOURCE | SOURCE | IDENTIFIER |
| --- | --- | --- |
| Antibodies |  |  |
| Bacterial and Virus Strains |  |  |
| *E. coli*: Strain OP50 | *Caenorhabditis* Genetics Center | OP50 |
| Chemicals, Peptides, and Recombinant Proteins |  |  |
| D-(-)-Sorbitol | VWR | Catalog #97062-204 |
| Critical Commercial Assays |  |  |
| Deposited Data |  |  |
| Experimental Models: Organisms/Strains |  |  |
| *C. elegans*: Wild-type Bristol N2 | *Caenorhabditis*<br>Genetics Center | N2 |
| *C. elegans*: *egl-1*(n487) | CGC | MT1082 |
| *C. elegans*: *flvEx196*(*cat-4::invertedCochr,myo-2::mcherry*);<br>*flvEx197*(*egl-6::Cre,myo-3::mcherry*) | This Paper | SWF502 |
| *C. elegans*: *egl-1*(n487); *flvEx196*(*cat-4::invertedCochr,myo-2::mcherry*); *flvEx197*(*egl-6::Cre,myo-3::mcherry*) | This Paper | SWF632 |
| *C. elegans*: *flvEx288*[*nlp-3::Gcamp5A (80ng/ul);nlp-3::mscarlet(20ng/ul)*]; *lite-1*(ce314) | This Paper | SWF640 |
| *C. elegans*: *flvEx304*[*sto-3::nlsGcamp7f(8ng/ul),tdc-1::nlsGcamp7f(10ng/ul), cat4prom68::GCaMP7b(50ng/ul),nlp-3::mscarlet(10ng/ul)*]; *lite-1*(ce314); *gur-3*(ok2245) | This Paper | SWF691 |
| *C. elegans*: *tph-1*(mg280); *flvEx196*(*cat-4::invertedCochr,myo-2::mcherry*); *flvEx197*(*egl-6::Cre,myo-3::mcherry*) | This Paper | SWF641 |
| *C. elegans*: *nlp-3*(n4897); *flvEx196*(*cat-4::invertedCochr,myo-2::mcherry*); *flvEx197*(*egl-6::Cre,myo-3::mcherry*) | This Paper | SWF915 |
| *C. elegans*: *tph-1*(mg280); *nlp-3*(n4897); *flvEx196*(*cat-4::invertedCochr,myo-2::mcherry*); *flvEx197*(*egl-6::Cre,myo-3::mcherry*) | This Paper | SWF873 |
| *C. elegans*: *flvEx196*(*cat-4::invertedCochr,myo-2::mcherry*);<br>*flvEx197*(*egl-6::Cre,myo-3::mcherry*);*unc-17*(*syb5779 syb5987*) | This Paper | SWF1012 |
| *C. elegans*: *egl-21*(n476); *flvEx196*(*cat-4::invertedCochr,myo-2::mcherry*); *flvEx197*(*egl-6::Cre,myo-3::mcherry*) | This Paper | SWF680 |
| *C. elegans*: *HSN::CoChR*; *HSN specific egl-21 KO*: *egl-21*(n476); *kySi61*[*loxP-egl-21genomic-loxP*]; *flvEx196*(*cat-4::invertedCochr,myo-2::mcherry*); *flvEx197*(*egl-6::Cre,myo-3::mcherry*) | This Paper | SWF717 |
| *C. elegans*: flp-2(gk1039); flvEx196(cat-4::invertedCochr,myo-2::mcherry);flvEx197(egl-6::Cre,myo-3::mcherry) | This Paper | SWF986 |
| *C. elegans*: flp-26(gk3015; flvEx196(cat-4::invertedCochr,myo-2::mcherry); flvEx197(egl-6::Cre,myo-3::mcherry) | This Paper | SWF838 |
| *C. elegans*: flp28(flv11); flvEx196(cat-4::invertedCochr,myo-2::mcherry); flvEx197(egl-6::Cre,myo-3::mcherry) | This Paper | SWF984 |
| *C. elegans*: flp-2(flv15); flp28(flv11); flvEx196(cat-4::invertedCochr,myo-2::mcherry); flvEx197(egl-6::Cre,myo-3::mcherry) | This Paper | SWF985 |
| *C. elegans*: flvEx404(Pegl6::HisCl; myo-2p::mcherry) | This Paper | SWF863 |
| *C. elegans*: flvls2[tph-1(NSM-specific fragment)::Chrimson, elt-2::mCherry] | Dag et al. (2023) | SWF117 |
| *C. elegans*: tph-1 (mg280); flvls2[Ptph-1(NSM-specific fragment)::Chrimson, Pelt-2::mCherry] | Dag et al. (2023) | SWF149 |
| *C. elegans*: tph-1(mg280); flvls2[tph-1(NSM-specific fragment)::Chrimson, elt-2::mCherry]; flvEx401[pegl-6::tph-1 cDNA, pmyo-2::mcherry] | This Paper | SWF855 |
| *C. elegans*: tph-1(mg280); flvls2[tph-1(NSM-specific fragment)::Chrimson, elt-2::mCherry]; flvEx495[ptph-1::tph-1 cDNA, pmyo-2::mcherry] | This Paper | SWF1011 |
| *C. elegans*: flvls2[tph-1(NSM-specific fragment)::Chrimson, elt-2::mCherry]; egl-1(n487) | This Paper | SWF831 |
| *C. elegans*: flvls2[tph-1(NSM-specific fragment)::Chrimson, elt-2::mCherry]; mod-5(n822) | This Paper | SWF854 |
| *C. elegans*: flvls2[tph-1(NSM-specific fragment)::Chrimson, elt-2::mCherry]; egl-1(n487); mod-5(n822) | This Paper | SWF853 |
| *C. elegans*: cat-1(syb7239); flvEx478[pegl-6::gfp1-10;pmyo-2::mcherry] | This Paper | SWF991 |
| *C. elegans*: flvEx201[tag168::mscarlet(5ng/ul); cat4prom68::gfp PCR (10ng/ul)] | This Paper | SWF506 |
| *C. elegans*: tax-2(p691) | CGC | PR691 |
| *C. elegans*: ocr-2(ak47) | CGC | CX4535 |
| *C. elegans*: tax-4 (p678) | CGC | CX13078 |
| *C. elegans*: -AWB: pels1715 [str-1p::mCasp-1 + unc-122p::GFP] | CGC | JN1715 |
| *C. elegans*: -AQR/PQR/URX: qals2241[gcy-36::egl-1; gcy-35::GFP; lin-15(+)] | CGC | CX7102 |
| *C. elegans*: -BAG: *kyls536 (flp-17p::p17 domain of human Caspase3::sl2::gfp; elt-2::GFP)*; *kyls538 (glb-5p::p12 domain of human Caspase3::sl2::gfp; elt-2::mcherry)* | CGC | CX11697 |
| *C. elegans*: -ASI: *oyls84(gpa-4p::TU#813+gcy-27p::TU#814+gcy-27p::GFP+unc-122p::DsRed)* TU#813 and TU#814 are split caspase vectors | CGC | PY7505 |
| *C. elegans*: -ASJ: *mglS40([daf-28p::nls-GFP]; jxEx100[trx-1::ICE + ofm-1::gfp])* | CGC | ZD762 |
| *C. elegans*: -ASK: *qrIs2[sra-9::mCasp1]* | CGC | PS6025 |
| *C. elegans*: -AFD: *ttx-1(p767)* | CGC | PR767 |
| *C. elegans*: -AIA: *kyEX4745[gcy-28dp::unc-103(gf)::sl2::mCherry, elt-2::mCherry]* | CGC | CX14597 |
| *C. elegans*: -AWC, ASE: *ceh-36(ky640)*; *kyls140 l[Pstr-2::GFP, lin-15(+)]* | CGC | CX5922 |
| *C. elegans*: -AIB: *flvEx356[inx-1::unc-103::sl2GFP(30ng/uL); myo-2::mCherry (5ng/uL)]* | This Paper | SWF763 |
| *C. elegans*: -ASK, ASI: *oyls84(gpa-4p::TU#813+gcy-27p::TU#814+gcy-27p::GFP+unc-122p::DsRed)* ;*qrIs2[sra-9::mCasp1]* | This Paper | SWF455 |
| *C. elegans*: *flvEx243[tax-4::tax-4 (50ng/ul), elt-2::gfp (5ng/ul)]* ;*tax-4 (p678)* | This Paper | SWF566 |
| *C. elegans*: *flvEx275[gcy-33::tax-4 (20ng/ul), elt-2::gfp (5ng/ul)]* ;*tax-4 (p678)* | This Paper | SWF615 |
| *C. elegans*: *flvEx497[sra-9::tax-4 (50ng/ul),elt-2::gfp (5ng/ul)]* ;*tax-4 (p678)* | This Paper | SWF1014 |
| *C. elegans*: *flvEx496[srg-47::tax-4 (50ng/ul),elt-2::gfp (5ng/ul)]* ;*tax-4 (p678)* | This Paper | SWF1013 |
| *C. elegans*: *flvEx298[srh-11::tax-4 (50ng/ul),elt-2::gfp (5ng/ul)]* ;*tax-4 (p678)* | This Paper | SWF674 |
| *C. elegans*: *flvEx285[srg-47::tax-4 (30ng/ul), sra-9::tax-4 (30ng/ul),srh-11::tax-4 (30ng/ul),elt-2::gfp (5ng/ul)]* ;*tax-4 (p678)* | This Paper | SWF634 |
| *C. elegans*: *flp-17(n4894)* | CGC | MT15933 |
| *C. elegans*: *flp-10(ok2624)* | CGC | RB1989 |
| Software and Algorithms |  |  |
| MATLAB (2021) | Mathworks | https://www.mathworks.com |
| Streampix (v7.0) | Norpix | https://www.norpix.com |
| Adobe Illustrator | Adobe | https://www.adobe.com |
| Other |  |  |
| SP-20000M-USB3 CMOS camera | JAI | N/A |
| Micro-NIKKOR 55mm f/2.8 lens | Nikon | N/A |
| 10x25 White Panel LED backlight, 24VDC | Metaphase Technologies | Cat#MS-BL10X25-W-24-ILD-PS |
| Precision LED Spot Light, 625nm, 40W, Type H | Mightex | Cat#BLS-PLS-0625-030-40-S |
| BioLED Light Source Control Module | Mightex | Cat#BLS-13000-1 |

### Plasmids and Promoters

Plasmid backbones: *C. elegans* codon-optimized GCaMP7b and GCaMP7f open reading frames were synthesized and inserted into the pSM vector. The intersectional promoters, consisting of the inverted/floxed vector and the Cre vector were previously described (Flavell et al., 2013). The HisCl1 plasmid was previously described (Pokala et al., 2014). For *tax-4* rescue, we used the previously described *tax-4* cDNA (Macosko et al., 2009), but moved it into a pSM-t2a-GFP vector backbone for expression. For *tph-1* rescue, we used a previously described cDNA (Zhang et al., 2005), which we moved into the pSM vector.

Promoters used in this study: *egl-6* (Flavell et al., 2013), *cat-4* (full length, 4kB immediately upstream of *cat-4* start codon), *cat-4prom68* (Lloret-Fernández et al., 2018), *sto-3* (Ji et al., 2021). The NSM-specific promoter was a 158 bp fragment of *tph-1* promoter, validated to be NSM-specific in our previous work through GFP expression and Ribotagging analysis (Rhoades et al., 2019). In addition, we showed that the resulting NSM::Chrimson line used here has no light-induced egg-laying even at maximum light intensities tested (Dag et al., 2023), further confirming that it confers no HSN-specific expression.

### New alleles generated in this study

The *egl-21* cell-specific deletion strain was constructed in an *egl-21(n476)* mutant background. For the mosSCI insertion, the *egl-21* genomic region (spanning entire genomic region up to adjacent genes in both directions) was inserted into the mosSCI insertion site on chromosome IV (Frøkjaer-Jensen et al., 2008). LoxP sites that we inserted into the *egl-21* single-copy rescue allow for Cre-dependent deletion of exons 2 through 5 of the *egl-21* gene, which is the majority of the coding sequence.

The conditional knockout allele of *unc-17* was constructed via iterative rounds of CRISPR/Cas9 genome editing. One loxP site was inserted ~250bp after the end of the *unc-17* 3’UTR. Another loxP site was inserted in the intron before the last coding exon (which encodes the majority of the UNC-17 protein). As described in the text, we found that pan-neuronal Cre expression in this strain gave rise to animals with an Unc phenotype, validating that the loxP sites work effectively.

The cell-specific fluorescent labeling strain for *cat-1* was generated via CRISPR/Cas9 genome editing. Three tandem repeats of the GFP11 sequence separated by short linker sequences (gly-gly-ser-gly-gly) were inserted immediately before the *cat-1* stop codon.

### Multi-animal behavioral recordings

Multi-animal recordings of *C. elegans* locomotion were conducted as previously described (Rhoades et al., 2019). One day old adult animals of the indicated genotypes were transferred to NGM plates with or without OP50 bacteria. For animals that were fasted, animals were transferred to NGM plates without OP50 for three hours prior to recording. All animals were recorded using Streampix software at 3 fps. JAI SP-20000M-USB3 CMOS cameras (5120×3840, mono) with Nikon Micro-NIKKOR 55mm f/2.8 were used. White-panel LEDs (Metaphase) provided backlighting. Videos were analyzed using previously-described custom MATLAB scripts (Rhoades et al., 2019). For optogenetic stimulation, light was supplied from a 470nm (for CoChR; 0.5 mW/mm2) or 625nm (for Chrimson; 0.6 mW/mm2 unless otherwise specified) Mightex LED at defined times in the video.

### Single-animal behavioral recordings

For joint recordings of egg-laying and locomotion, we used previously described custom-built single worm tracking microscopes (Cermak et al., 2020). These custom microscopes have a live-tracking function that permits long-term recording of single moving animals. Animals recorded on these microscopes were transferred to 30% peptone NGM plates with OP50 prior to recording (the reduced peptone ensured that bacterial lawns would remain thin, which is a requirement for the live-tracking function on the microscopes). LabView software controlled the microscope and acquired the images. Data were then analyzed in R Studio and MATLAB. For optogenetic stimulation, 532nm laser light was supplied at defined times at an intensity of 250 uW/mm^2^.

### HSN Axotomy

Laser axotomy was performed as previously described (Byrne et al., 2011) with a few modifications. L4 stage transgenic animals were transferred to a 3% agarose pad and immobilized with 2.5mM levamisole in M9 buffer. Animals were visualized with a Nikon Eclipse 80i microscope, 100x Plan Apo VC lens (1.4 NA), Andor Zyla sCMOS camera and a Leica EL6000 light source. HSN axons were severed anterior to the vulval presynaptic region and before they extend to the ventral nerve cord using a 435nm nitrogen pulsed MicroPoint laser fired at 20 Hz. Both HSNL and HSNR axons were sequentially severed by gently rolling the animal from one side to the other after the first axotomy. To facilitate rolling, grooved agarose pads were stamped with a portion of a vinyl record (Rivera Gomez and Schvarzstein, 2018). ‘Mock’ control animals underwent the same immobilization and rolling protocol but were not axotomized. Both mock and axotomized animals were immediately recovered in M9 buffer and transferred to OP50 seeded NGM plates for behavioral analyses 20 hours later.

### Freely-moving HSN Calcium Imaging

HSN calcium imaging in freely-moving animals was conducted as previously described (Rhoades et al., 2019) with a few small modifications. Animals were mounted on flat agar, with freshly seeded OP50 bacterial food. They were enclosed in a small chamber with a rubber gasket and cover glass and GCaMP/mCherry data was recorded at 20 fps (with 10ms exposure times). Imaging was performed with a 4x/0.2NA objective and data was acquired on two Andor Zyla 4.2 Plus sCMOS cameras. A Cairn TwinCam beam splitter was used to separate GCaMP and mCherry signals. The GCaMP/mCherry imaging at 20 fps was interleaved with 2 fps brightfield imaging, achieved via NI-DAQ triggering of different light sources using the NIS Elements Illumination Sequence module.

For data analysis, the HSN soma was tracked (using the bright mCherry signal) using custom ImageJ macros. After cell positions were determined by ImageJ tracking, the GCaMP and mCherry signals were extracted from each ROI at each time point. Background was subtracted from each signal and then the ratio of the two background-subtracted fluorescence measurements was taken. Time points of egg-laying were manually determined from the brightfield images.

### Immobilized Calcium Imaging

Calcium imaging of HSN and RIB, as well as different HSN compartments, in immobilized animals was conducted on a previously described spinning disk confocal microscope (Ji et al., 2021). Animals were mounted on 5% agar with 0.05um beads for immobilization (Kim et al., 2013). They were imaged using a 20X/0.95 objective coupled to a 5000 rpm Yokogawa CSU-X1 spinning disk unit with a Borealis upgrade. Z-stacks were collected with NIS Elements software. For data analysis, data were converted to maximum intensity projections (RIB and HSN were typically in different z-planes), ROIs were manually drawn around the somas of RIB and HSN, and then background-subtracted intensities within the ROIs were calculated.

### Egg Counting Assays and Osmolarity Exposure

Sorbitol was used to increase the osmolarity in the NGM plates to desired osmolarity (300 mOsm −450 mOSm), and added together with CaCl_2_ and KPO_4_ buffer to the NGM agar. Assay plates were seeded with 200 ul OP50 per plate, and lids were left open to allow OP50 to be dried before putting away for the assay. L4s animals were picked to OP50-seeded NGM plates the day before the assays, and grown for 20-24 hours to become gravid adults. On the day of the experiment, adult animals were transferred onto seeded control or high osmolarity plates (10 animals per plate), and were left to lay eggs for an hour. After an hour, animals were removed from assay plates and the number of eggs laid was counted. The percentage of egg laid was calculated by the number of eggs laid on high osmolarity plates divided by the number of eggs laid on control plates.

### Exploration assays

Exploration assays that provide a reliable measure of roaming versus dwelling behavior were performed as previously described (Flavell et al., 2013) with minor modifications. Single L4 animals were picked to NGM plates with OP50 bacteria seeded 1 day prior. They were then left to explore the plates for 16 hours, after which the plate was superimposed on a transparent grid and the number grid squares that the animal tracks traversed was quantified. In some experiments (as indicated in the text), animals were mounted as adults on plates and allowed to explore for 3 hours, and then the number of squares was counted.

---

**Figure S1, Related to Figure 1.**
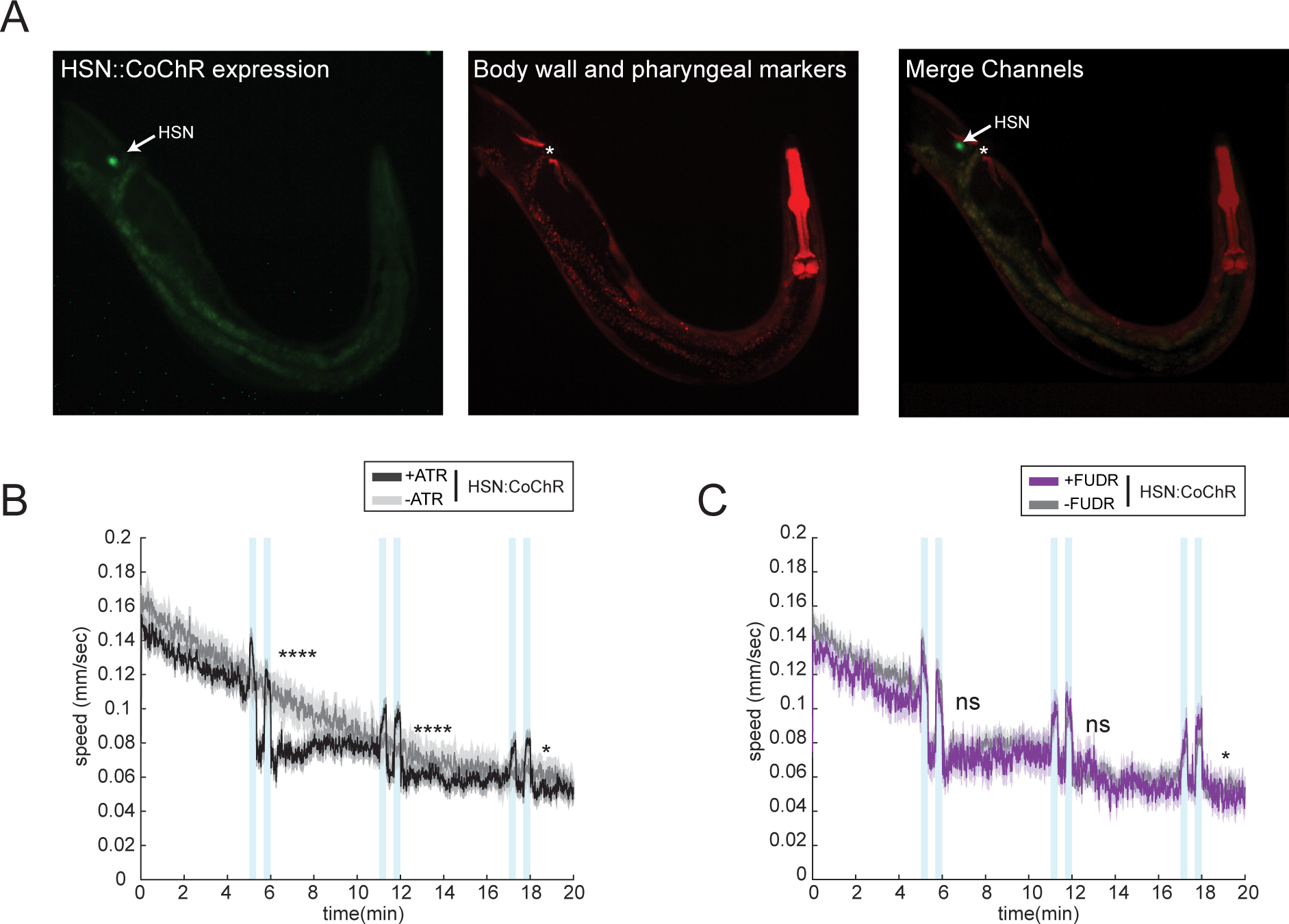
(A) Representative images of HSN::CoChR-sl2-GFP expression (left), along with body wall and pharyngeal markers (middle), and a composite image (right). Arrows indicate HSN soma. Asterisks indicate vulva position. (B) Full recordings of animal speed during several bouts of HSN::CoChR stimulation for animals recently transferred to plates with food. Data are shown for HSN::CoChR animals with and without ATR. N= 195 animals for the ATR group, and 159 animals for the no-ATR control. *p<0.05, ****p<0.0001, Bonferroni-corrected t-test. (C) Animal speed during several bouts of HSN::CoChR stimulation for animals recently transferred to food plates. Data are shown for HSN::CoChR animals either treated with FUDR or not. N= 69 animals for the FUDR-treated group, and 195 animals for the no-FUDR control. *p<0.01, Bonferroni-corrected t-test.

**Figure S2, Related to Figure 2.**
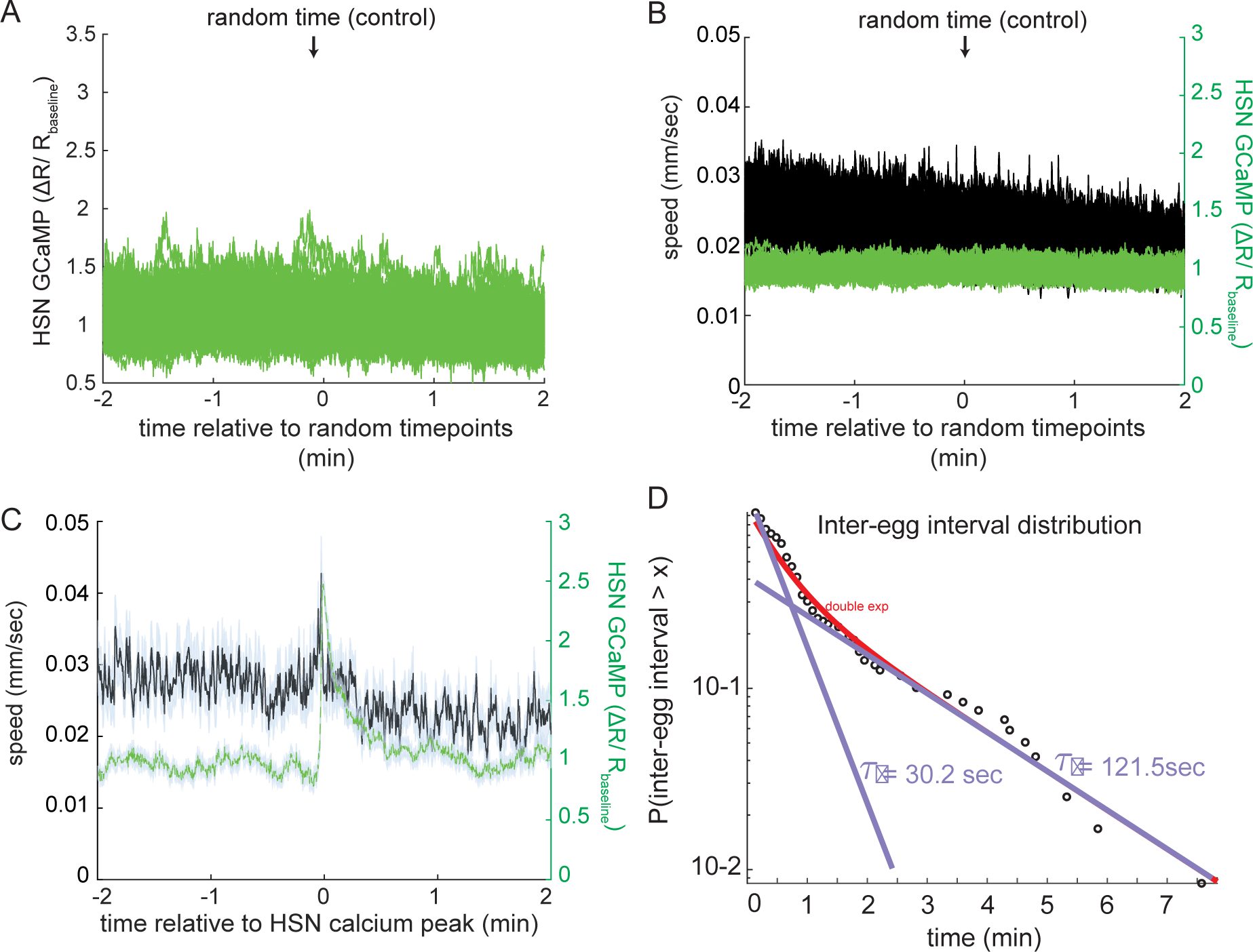
(A) Event-triggered averages showing average HSN GCaMP signal surrounding randomly chosen timepoints, as a control for Fig. 1B. Each line is the mean of 16 timepoints (matching the N in Fig. 2B), and this control was run 100 times, resulting in 100 lines. (B) Event-triggered averages showing average speed (black) and HSN GCaMP (green) surrounding randomly chosen timepoints, as a control for Fig. 2C. Each line is the mean of 104 timepoints (matching the N in Fig. 2C), and this control was run 100 times, resulting in 100 lines. (C) Event-triggered average showing average animal speed surrounding HSN calcium peaks. This plot only includes HSN calcium peaks that were not accompanied by egg-laying. Note that speeding still occurs during these peaks. (D) Complementary cumulative distribution function (ccdf) showing distribution of intervals between HSN calcium peaks. This distribution was best fit by a double exponential (red). The slopes of each exponential are shown in blue and the tau values are also displayed. The shorter distribution is characterized by a tau of ~30s, whereas the longer distribution is characterized by a tau of ~2min.

**Figure S3, Related to Figure 3.**
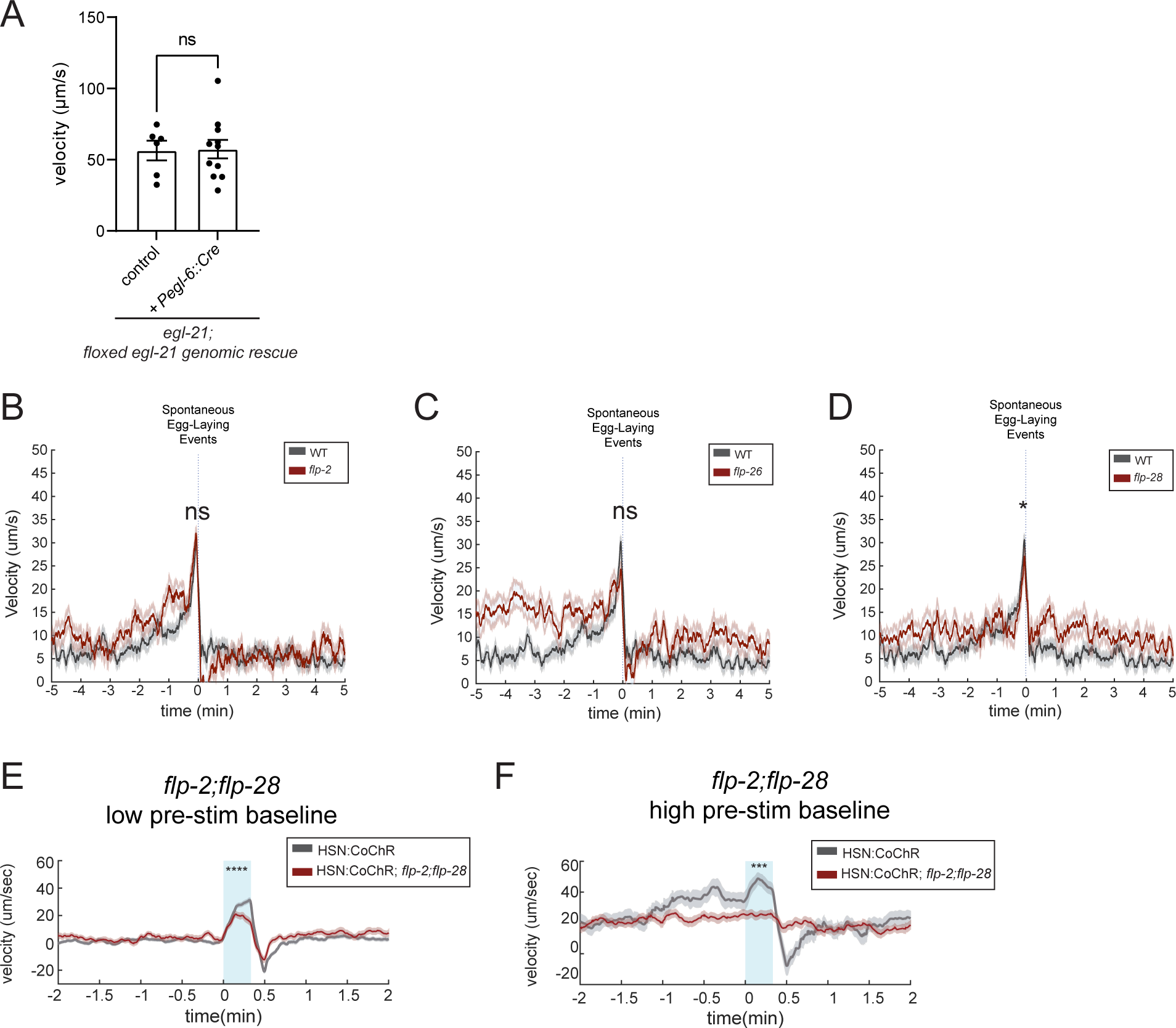
(A) Baseline mean velocity of *egl-21; floxed egl-21 genomic rescue* animals without (control) or with *pegl-6::Cre* expression in the absence of food. Dots show individual animals; bars are means and error bars are SEM. N= 6 animals for control and 11 animals for *pegl-6::Cre* expressing animals. (B) Event-triggered averages for time periods surrounding native egg-laying events in *flp-2(gk1039)* animals. N= 194 egg-laying events across 6 animals. (C) Event-triggered averages for time periods surrounding native egg-laying events in *flp-26(gk3015)* animals. N= 162 egg-laying events across 6 animals. (D) Event-triggered averages for time periods surrounding native egg-laying events in *flp-28(flv11)* animals. N= 159 egg-laying events across 6 animals. *p<0.05, unpaired t-test. (E and F) Event-triggered averages showing average changes in velocity in *flp-2(flv15);flp-28(flv11)* animals in response to HSN::CoChR activation via light illumination. Gray provides data from WT animals as a control and reference. Lines show means and error shading shows SEM. (D): Data with animals dwelling at low velocity (<20µm/sec) before light illumination. N =249 stimulation events for WT and 129 stimulation events for *flp-2(flv15);flp-28(flv11)* animals. (E): Data with animals having higher velocity baseline (>=20µm/sec) before light illumination. N =21 stimulation events for WT and 46 stimulation events for *flp-2(flv15);flp-28(flv11)* animals. ****p<0.0001, unpaired t-test.

**Figure S4, Related to Figure 4.**
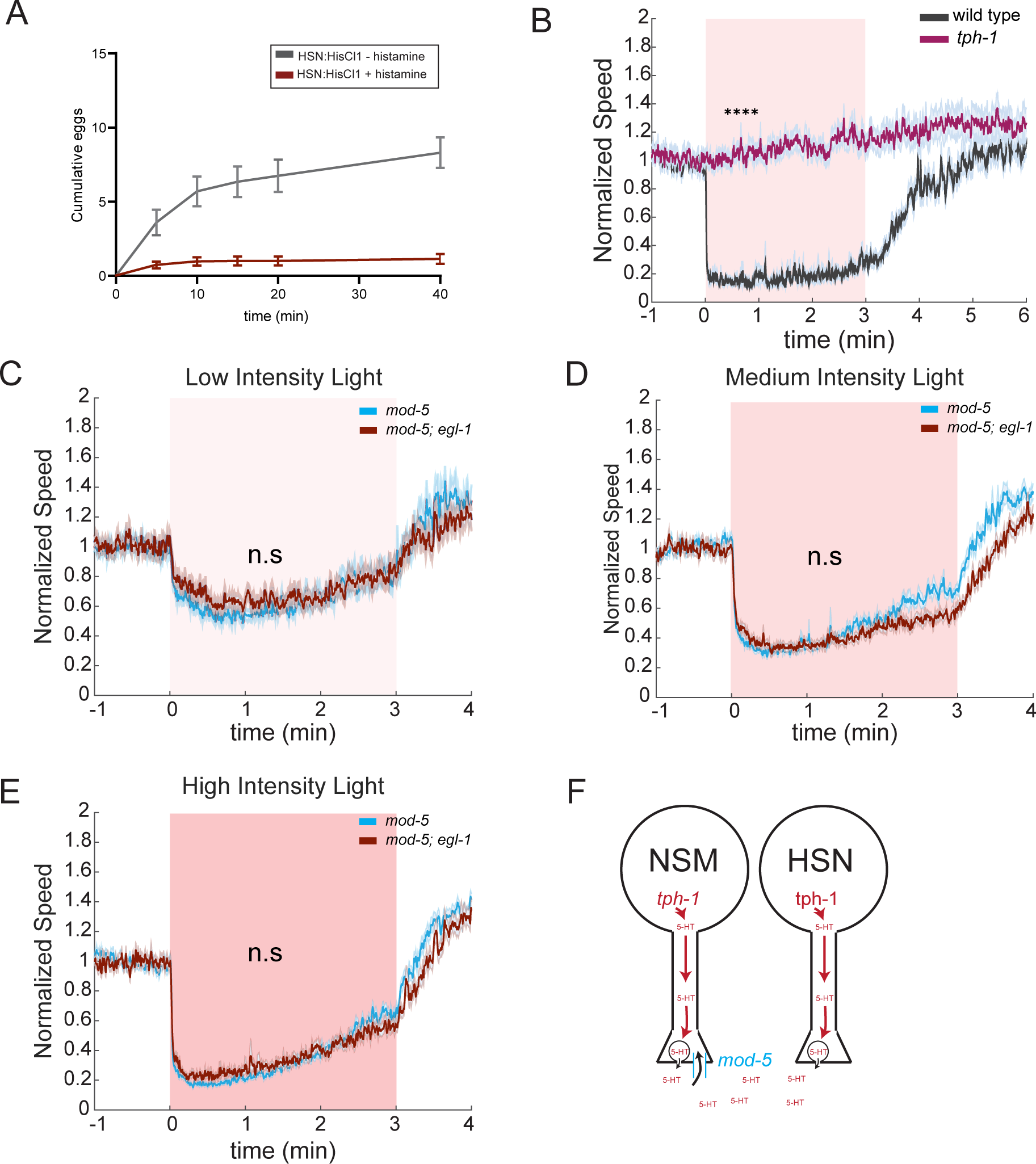
(A) Egg-laying of HSN::HisCl animals either exposed to histamine or not. Animals for this experiment were transferred to +his or −his plates immediately before this assay (i.e. at t=0min), and eggs laid were counted at different time points, up to 40min after transfer. Note that egg-laying is reduced even at the first time point, indicating the HSN::HisCl inhibits egg-laying within minutes of first exposure to histamine. N=35 plates for HSN::HisCl with histamine and 20 plates for no histamine control group, with 3 animals on each plates. (B) Event-triggered averages depicting the average change in animal speed upon NSM::Chrimson stimulation with red light illumination in wild-type and *tph-1(mg280)* animals. Animals were starved for 3 hours before the assays. Lines show means; and error shading shows SEM. N= 77-390 animals. ***p<0.0001, unpaired t-test. (C to E) Event-triggered averages depicting the average change in animal speed upon NSM::Chrimson stimulation with different light intensity in *mod-5(n822)* and *mod-5(n822);egl-1(n487gf)* animals, with panel C being the lowest intensity and panel E the highest. N= 51-248 animals. (F) Cartoon illustrating serotonin release and re-uptake by NSM and HSN neurons.

## Notes

### Competing Interest Statement

The authors have declared no competing interest.

